# The role of contact guidance and ECM remodelling in cancer invasion: a computational study

**DOI:** 10.1101/2025.09.02.673661

**Authors:** Margherita Botticelli, John Metzcar, Paul Macklin, Pradeep Keshavanarayana, Fabian Spill

## Abstract

The extracellular matrix (ECM) is a complex network of fibrous proteins and other macromolecules that provides both structural support and directional cues that regulate cancer cell invasion and tumour progression. Its fibre organisation plays a critical role in directing migration through contact guidance, while cancer cells simultaneously remodel the matrix through biochemical and mechanical interactions. However, the interplay between ECM architecture, chemical gradients, and matrix remodelling remains poorly understood. Mathematical modelling offers a powerful approach to explore how ECM architecture regulates this process. We present a hybrid computational model, implemented in PhysiCell, that integrates a discrete agent-based cell model with a continuous representation of the chemical microenvironment and ECM microstructure. In an advance over previous efforts, we adapt several mechanisms to a fibre-focused representation of the ECM, including “cell-front” ECM sensing reflecting protrusion-driven engagement of the matrix, contact-guided cell movement via ECM fibre orientation integrated with chemotaxis bias, and proliferation regulated by oxygen and mechanical pressure. In addition, we introduce a new mechanical mechanism for ECM density displacement alongside degradation to simulate how cells redistribute matrix fibres. Simulations reveal how the interplay between initial fibre alignment, anisotropy (fibre-fibre alignment correlation), and fibre reorientation capacity affects invasion, and how competing mechanical and chemical cues influence the invasive potential of tumour cells. Furthermore, our results demonstrate that the balance between degradation and displacement strongly affects invasion dynamics, with high displacement promoting the formation of dense ECM rims around tumour spheroids, whereas increased degradation enables greater invasive spread. This work provides mechanistic insights into bidirectional interactions between cancer cells and the surrounding ECM, highlighting how structural remodelling of the ECM influences the invasive potential of cancer cells.

**Author summary:** Cancer cells invade surrounding tissue by interacting with the extracellular matrix (ECM), a fibrous network of proteins that provides both mechanical support and directional cues for migration. Experiments have shown that the orientation of ECM fibres can either promote or hinder invasion, but it remains difficult to disentangle the underlying mechanisms because cells both respond to and actively remodel the matrix. To address this challenge, we developed a computational model that simulates how cancer cells migrate through and reshape the ECM. The model combines individual cell behaviour with a representation of ECM structure, including fibre orientation, anisotropy (fibre-fibre alignment correlation), and density. Our simulations show that invasion depends on the initial fibre orientation and on chemical cues. We also introduce a mechanism that allows cells to mechanically displace the matrix, revealing how the balance between matrix degradation and physical pushing can generate either compact tumour growth or sparse invasion. These results help explain how physical interactions between cells and their environment shape tumour invasion. More broadly, the framework provides a tool to explore how mechanical and chemical signals together regulate collective cell migration in cancer and other biological systems.

## 1 Introduction

The extracellular matrix (ECM) is a structurally and functionally complex network of macro-molecules, predominantly composed of fibrous proteins (*e*.*g*., collagen), that play a crucial role in tumour growth and invasion ^35^. Its structural, mechanical and biochemical properties critically regulate cellular processes including differentiation, proliferation, and migration ^15;27^. Through a process called mechanosensing, the cells can interpret and respond to mechanical cues from the ECM, including matrix density, stiffness, porosity, viscosity and fibre orientation, which in turn influence their behaviour ^7;27;14^. Cancer cells actively remodel the ECM through various mechanisms, including enzymatic degradation, deposition of new matrix proteins, and mechanical reorientation of existing fibres, thereby creating a favourable environment that facilitates their migration and invasion ^34;4^. Aligned ECM fibres can guide cell migration, facilitating directional invasion, while isotropic matrices may inhibit coordinated movement, as reduced fibre alignment has been shown to diminish migratory persistence and speed ^31^. This fibre-guided migration, often referred to as contact guidance, is increasingly recognised as a central mechanism underlying collective invasion and metastatic potential^21;34^.

Understanding these complex bidirectional interactions between the cancer cells and their surrounding environment is crucial for deciphering tumour progression and designing effective interventions. In-vitro models are widely utilised to investigate tumour development and invasion within controlled three-dimensional (3D) environments ^24^. However, purely experimental investigation of these interactions is often difficult, given their complexity and multiscale nature. Computational models have demonstrated to be invaluable tools for simplifying these complexities and predicting cancer dynamics. These models employ various approaches, including continuum, discrete (*e*.*g*., agent-based), and hybrid models, each suited to capture different aspects of cancer cell behaviour. Hybrid models, in particular, offer a significant advantage by representing multi-scale interactions between cancer cells and the ECM, coupling individual cell behaviours with macroscopic tissue-level dynamics ^12;3^.

Numerous computational models have been developed to investigate the interplay between cells and the fibrous ECM^12;3;2;13^. Recent notable examples include the work by Noël et al. ^16^, who developed PhysiMeSS, an add-on to the agent-based modelling framework PhysiCell ^6^, designed to model ECM fibres as discrete agents. Each ECM fibre agent is represented by individual 2D segments or 3D cylinders, with properties such as stiffness. This formulation enables the simulation of physical interactions between cells and other fibres, as well as fibre alignment and degradation. Fibre remodelling and contact guidance play an important role in wound healing. For instance, Richardson and Holmes ^22^ investigated contact guidance and fibre reorientation using a finite-element model in the context of cardiac wound healing. Some studies focus more on the ECM’s role in cancer invasion specifically. Poonja et al. ^19^ built a novel multi-cellular lattice-free agent-based model that captures the dynamic interplay between cell-generated forces and ECM architecture, reproducing transitions between Tumour-Associated Collagen Signatures (TACS), which are correlated with the invasive potential of tumours. Using a single ECM-density field, Ruscone et al. ^25^ explored cancer cell invasion using PhysiBoSS ^10;18^, a framework coupling Boolean network simulation utilising the Boolean network simulator MaBoSS ^29^ and the ABM framework PhysiCell ^6^. Sander ^26^ provided another example focused on cancer invasion, in this case using a deterministic, anisotropic tensor-based approach. While some existing models capture both contact guidance and the influence of physical ECM fibres on cell behaviour, they do so either with highly detailed fibre-level simulations or coarse bulk-density approaches. None have employed a meso-scale framework that balances computational efficiency with sufficient structural detail to study how ECM architecture guides cell migration and invasion.

We significantly advance our previous computational model ^1^ of density-dependent spheroid growth by incorporating a more nuanced representation of microenvironmental dynamics. While our earlier work focused on ribose-induced stiffening, we now introduce several biophysical mechanisms to capture the complexity of the ECM microstructure. First, we integrate ECM fibre orientation and anisotropy to guide cell migration. Next, we enhance our “cell-front” framework by introducing a mechanical function for ECM density displacement alongside degradation. Finally, we implement a revised proliferation mechanism whereby the cell division is now explicitly regulated by oxygen concentration and mechanical pressure. These innovations, combined with established features such as the chemical microenvironment, cell-cell adhesion and repulsion and chemotaxis already integrated with PhysiCell ^6^, and ECM remodelling (fibre reorientation, anisotropy increase, and density degradation) ^13^, create a more biologically-grounded framework for investigating how ECM architecture, anisotropy, and reorientation capacity influence invasive cell behaviour. This model enables a systematic study of how initial ECM configurations and their remodelling over time affect cancer cell growth and invasion, and provides theoretical insights into key experimental observations ^20;30^, reinforcing the biological relevance of our approach.

## 2 Methods

### 2.1 General framework

We present a hybrid discrete-continuous model developed in PhysiCell ^6^, extending our previous work ^1^. PhysiCell is an open-source, multiscale modelling framework that simulates multicellular biological systems, representing cells through an off-lattice, agent-based model ^12^ coupled with a continuous model for the surrounding chemical microenvironment^6^. The simulation of diffusible chemical substrates, such as oxygen and other nutrients, is handled by BioFVM, PhysiCell’s efficient, parallelised multi-substrate diffusion solver ^5^. BioFVM simulates substrate dynamics through reaction-diffusion-decay partial differential equations, accounting for diffusion, decay, and cellular uptake/secretion.

We adopt the ECM framework of Metzcar et al. ^13^, which represents the ECM on a lattice of voxels, each characterised by average fibre orientation, anisotropy (average local fibre-fibre alignment correlation), and density (average local volume fraction of fibres).

In our previous work, we examined the impacts of ribose stiffening on spheroid growth and invasion using solely ECM density ^1^. Here, we extend that model by removing the ribose component and incorporating fibre orientation and anisotropy into the ECM representation. This allows us to investigate how initial fibre architecture and its remodelling influence cancer cell migration and proliferation, particularly through contact guidance and directional ECM sensing. While the earlier model focused primarily on how ECM density and stiffening regulate invasion speed and spheroid growth, the present work investigates how ECM architecture, specifically fibre orientation and anisotropy, guides invasion dynamics through contact guidance and cell-driven matrix remodelling.

Our model consists of four key parts including both pre-existing and novel functions:

- Chemical microenvironment: describes the spatiotemporal evolution of chemical substrates within the system.
- Cell motion: influenced by both cell-cell interactions and cell-microenvironment interactions, encompassing cell-ECM interactions and chemotaxis towards specific chemical substrates.
- ECM remodelling: describes active cellular modification of ECM microstructure (fibre orientation, anisotropy, and density).
- Cell proliferation: chemical substrates and mechanical pressure impact cell proliferation.

Three new mechanisms are introduced in this work compared to Botticelli et al. ^1^ and Metzcar et al. ^13^ . First, we define a revised motility direction function that integrates contact guidance with chemotaxis (Section 2.2). Second, we introduce an ECM density displacement mechanism (Section 2.3). This feature allows ECM density to be relocated into the voxel the cell is moving towards, simulating mechanical displacement of matrix fibres. A similar mechanism was proposed by Poonja et al. ^19^ to model fibril stiffening due to cell-generated forces during division. Finally, while in Botticelli et al. ^1^ inhibition of proliferation was based on ECM density and neighbour count, here we integrate recent work in Wang et al. ^32^, Rocha et al. ^23^, and Johnson et al. ^8^ where proliferation is regulated by oxygen concentration and by mechanical pressure from neighbouring cells (Section 2.4), capturing the combined effects of ECM repulsion and cellular overcrowding, offering a more physiologically relevant model for growth inhibition.

In this article, while ECM remodelling of fibre orientation, anisotropy and degradation follows the partial differential equations (PDEs) of Metzcar et al. ^13^ (S1 text Section 1.3), cell-microenvironment interactions reuse functions from our previous work ^1^ (S1 text Section 1.2), except for the novel motility direction function (Section 2.2). We utilise the existing Physicell frameworks for chemical microenvironment dynamics and cell-cell adhesion and repulsion without further modification (S1 text Section 1) . As in our earlier model ^1^, we employ a pseudo-2D (or constrained 3D) model to match *z*-slice experimental imaging. We also retain the “cell front” voxel selection rule for ECM interactions; discussed in details in S1 text Section 1.

### 2.2 Cell motility direction: ECM-guided and chemotactic migration

Cell migration in biological tissues typically arises from a protrusion-adhesion cycle in which cells extend protrusions (*e*.*g*. lamellipodia or filopodia), form transient adhesions with ECM fibres, and generate traction forces through actomyosin contraction. In the centre-based representation used here, these subcellular processes are not modelled explicitly. Instead, they are coarse-grained into an effective motility velocity whose magnitude depends on ECM density and whose direction is influenced by fibre orientation and chemotactic gradients.

While in Botticelli et al. ^1^, the cell’s motility direction was equal to a random unit vector (***d***_*i*,mot_ = ***d***_random_), here, building upon Metzcar et al. ^13^, we modified the direction of movement of the cell to be influenced by chemotaxis and fibre orientation as follows:

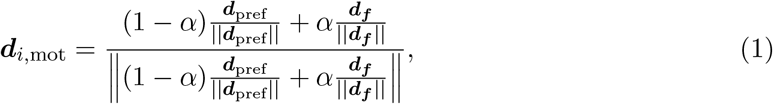

where *α* is the ECM sensitivity parameter. The ECM sensitivity parameter determines the extent to which a cell’s motility direction is governed by contact guidance along ECM fibres. In Equation (1), ***d***_pref_ is the “preferred direction” of movement, defined by a combination of random motion with chemotaxis bias. This combination is weighted by a chemotaxis bias parameter *β*, as:

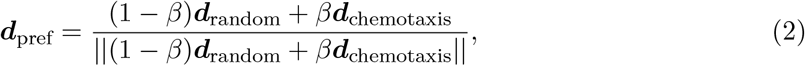

where ***d***_random_ is a unit vector with random direction sampled from a uniform distribution.

Finally, ***d***_***f***_ is the “fibre direction” of movement given by the fibre orientation ***f*** modulated by the anisotropy *a* which adds randomness to the vector. Since fibre orientation is independent of vector sign, we define the raw fibre direction as:

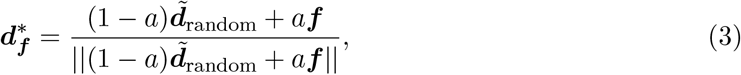

where 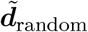 is a random unit vector generated similarly to ***d***_random_:

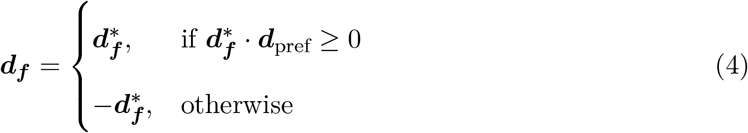

Higher anisotropy corresponds to higher fibre-fibre correlation, equivalent to having more parallel fibres, meaning there is less randomness in fibre orientation in a voxel.

### 2.3 ECM density displacement

In addition to the ECM degradation used in Botticelli et al. ^1^ and Metzcar et al. ^13^ (S1 text Section 1.3), we define the ECM displacement function. In the previous formulation, ECM remodelling occurred solely through degradation with ECM density target *ρ*_target_ = 0, corresponding to complete matrix removal by the cells. However, biologically, complete degradation to zero density is unrealistic. Hence, we set the degradation target (*ρ*_target_, Equation (16) in S1 text) to 0.5, corresponding to the ECM density at which individual cells reach maximum speed (Equation (13) in S1 text). Additionally, cells are more likely to remodel their surroundings by degrading the matrix to an intermediate density while displacing it, creating an optimal microenvironment that maintains both mechanical support and migratory efficiency. Accordingly, we present a new implementation where ECM degradation and displacement occur simultaneously. This approach balances local degradation with a displacement mechanism that redistributes the remaining ECM between neighbouring voxels. This process creates a depleted zone at the spheroid centre while progressively pushing the matrix outward, facilitating sustained invasion.

We assume all cells can relocate the ECM present in their current position’s voxel (voxel *A*, Figure 1) into the adjacent voxel in their direction of motion (voxel *B*, Figure 1) proportionally to their total speed. In this formulation, for simplicity, we assume that the ECM displacement is permanent. No elastic restoring force is included, and displaced ECM does not return to a reference configuration if cells move away or are removed. Thus, ECM displacement represents plastic remodelling of the matrix rather than reversible elastic deformation. Fibre orientation and anisotropy evolve exclusively through the reorientation and realignment dynamics defined previously in Metzcar et al. ^13^ (S1 text Section 1.3.) and are not directly altered by density displacement.

**Figure 1.**
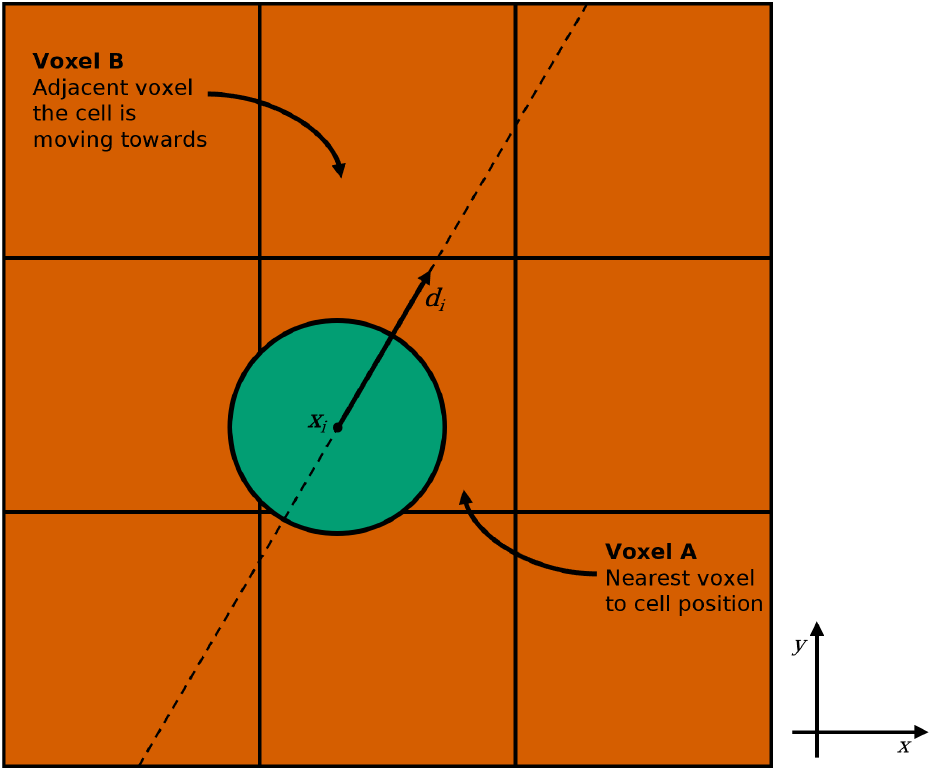
Schematic of ECM displacement. The green circle represents a cell located at position ***x***_*i*_, with direction of motion ***d***_*i*_. Voxel *A* is the nearest voxel to the cell’s current position, while voxel *B* is the adjacent voxel in the cell’s direction of motion. The dashed line represents the trajectory defined by the direction vector ***d***_*i*_, used to determine which voxel boundary the cell will intersect first.

Each voxel stores an ECM density *ρ ∈* [0, 1]. Let *ρ*_*A*_ and *ρ*_*B*_ denote the densities in voxels *A* and *B*, respectively, and let *r*_*d*,disp_ be the displacement rate and *s*_*i*_ the cell’s total speed. If voxel *A* is not empty (*ρ*_*A*_ *>* 0), the displacement updates are given by:

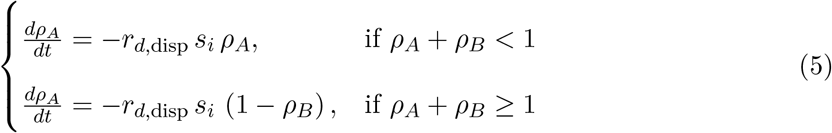

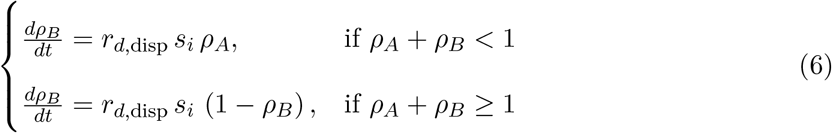

ensuring *ρ*_*A*_ and *ρ*_*B*_ *∈* [0, 1]. When *ρ*_*A*_ + *ρ*_*B*_ *<* 1, the transfer conserves mass between the voxels. When *ρ*_*A*_ + *ρ*_*B*_ *≥* 1 the transfer fills voxel *B* and reduces *A* by the corresponding amount so that *ρ*_*B*_ never exceeds 1.

To identify voxel *B*, we trace the line passing through the cell’s position ***x***_*i*_ with slope equal to its final direction of motion ***d***_*i*_ and find which adjacent voxel it intersects first, as shown in Figure 1.

### 2.4 Cell proliferation

Cell proliferation is a critical process in our model, simulated using the simplified “live” cell cycle model from PhysiCell ^6^, where the cell cycle is represented by a single “live” phase. The cell division rate is modulated by an inhibition of proliferation function (*f*_*IP*_), which accounts for the combined effects of nutrient concentration and pressure from neighbouring cells. The probability of a cell dividing within a time interval [*t, t* + Δ*t*] is given by:

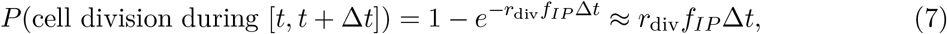

where *r*_div_ is the maximum division rate.

In our previous model in Botticelli et al. ^1^, the inhibition of proliferation function depended on ECM density and the number of neighbours. Instead, in this model, we take into account oxygen conditions and cellular pressure. The cell’s proliferative response to environmental oxygenation is modelled using a Hill function, as in Johnson et al. ^8^ . Specifically, we use a half-maximum of 21.5 mmHg and a Hill power of 4. The inhibition of proliferation due to pressure effectively captures the combined influence of ECM repulsion and cellular overcrowding. Although ECM density is not directly included in the definition of cell pressure, it indirectly influences it by constraining cell movement due to the repulsive forces between cells and the ECM (S1 text Section 1.2.3). In the case of a spheroid, the ECM surrounding the spheroid acts as a confining barrier. As cells proliferate and the spheroid expands, the cells at the boundary encounter resistance from the ECM. This resistance increases local cell packing and, consequently, the intercellular pressure.

Hence, we define the inhibition of proliferation function as

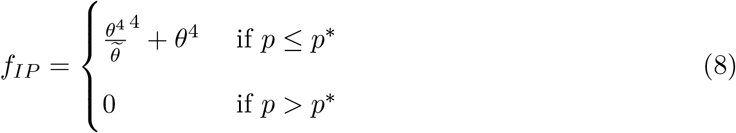

where *θ* is the current environmental oxygenation, 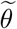 is the half-maximum, *p* is the current pressure experience by the cell, and *p*^*\**^ is the pressure threshold.

The pressure threshold, *p*^*\**^, is defined as the pressure a cell would experience when surrounded by six cells at their equilibrium distance, representing a packed 2D configuration. In PhysiCell, the pressure *p*_*i*_ on a cell *C*_*i*_ is calculated as:

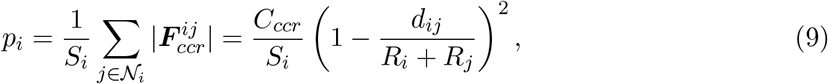

where *S*_*i*_ is the cell’s surface area, *𝒩*_*i*_ is the set of neighbouring cells, *C*_*ccr*_ is the cell-cell repulsion coefficient, *R*_*i*_ and *R*_*j*_ are the radii of the cells *C*_*i*_ and *C*_*j*_, respectively, and *d*_*ij*_ is the distance between their centres. This pressure is then non-dimensionalised by a scaling factor, 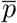, which represents the pressure a cell would experience in a 3D packing, surrounded by 12 cells. The non-dimensionalised pressure is given by:

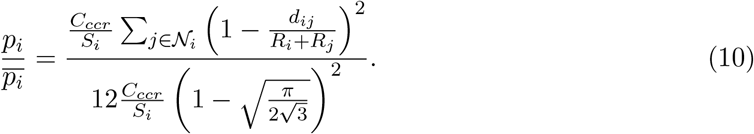

For a more detailed explanation of pressure calculation, refer to the PhysiCell documentation ^6^.

To determine the equilibrium distance, *d*, between two cells, we assume that the cell-cell adhesion and repulsion forces are balanced:

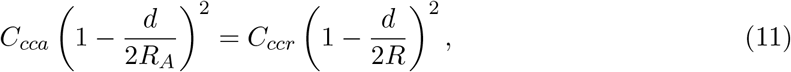

where *C*_*cca*_ is the cell-cell adhesion coefficient, *R*_*A*_ is the adhesion radius, and *R* is the cell radius.

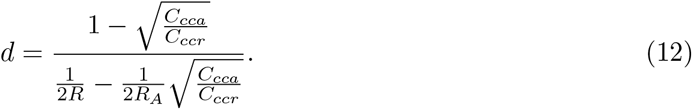

Finally, the pressure threshold *p*^*\**^ is defined using the equilibrium distance, considering the pressure exerted by six surrounding cells:

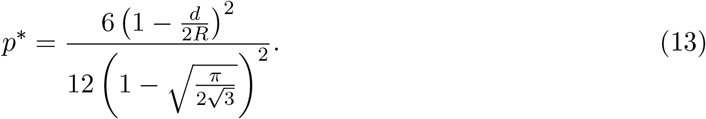

### 2.5 Metrics and computational details

To account for the stochastic nature of the model, we performed 10 replicates for each simulation. Across these replicates, we observed low standard deviations for all key metrics, which include normalised fibre orientation, cell spatial distribution, invasion, spheroid growth relative to *t* = 0, and Delaunay mean distance. These narrow distributions indicate a low relative error and confirm the consistency of our results. All simulations covered 96 hours, sampling every 60 minutes, and were performed in a domain of size 1000*×*1000*×*20 *µ*m^3^ to which cell movement is constrained (the domain boundaries act as impenetrable walls), with each ECM voxel of size 20*×*20*×*20 *µ*m^3^.

The methodologies for calculating spheroid growth relative to the initial time and Delaunay mean distance are consistent with our previous work ^1^. Spheroid growth measures the area covered by the cells with respect to the initial time, without over-counting overlapping regions, while Delaunay mean distance provides a measure of average cell-to-cell distance.

Normalised fibre orientation provides a measure of how aligned the ECM fibres are with the *y−*axis (Figure 1). It is calculated by first taking the fibre orientation angle for each voxel. Subsequently, the absolute difference between each average fibre angle and 90° is computed. These differences are then normalised by dividing by 90, resulting in values between 0 and 1. A value of 0 indicates that the fibre orientation in a voxel is perpendicular (aligned with the *y−*axis), while a value of 1 signifies a perfectly parallel (aligned with the *x−*axis) orientation. Finally, the median of all these normalised angles across the whole domain is taken to represent the overall normalised fibre orientation.

The cell spatial distribution metric quantifies how cells are positioned relative to a reference point in the simulation domain (origin). Distances from the origin are divided into equal-width bins of 20 *µ*m, and the number of cells within each bin is counted. The resulting distribution profiles reveal how far cells travel.

Invasion quantifies the extent to which cells migrate outwards. Specifically, for simulations depicting upward (or downward) invasion, we calculate the median of the distances of all cells’ *y*-position from their initial position (at the bottom in Section 3.1 or centre of the domain in Section 3.2). For spheroid-specific simulations (where the focus is on radial expansion), invasion is measured as the 95th percentile of the distances of all cells from the centre of the domain (Sections 3.3 and 3.4). This captures the outward spread of the most invasive cells.

Physicell was compiled using g++ (version 8.5.0), and post-processing analyses were conducted using Python (version 3.10.12). All simulations were performed on the BlueBEAR high-performance computing (HPC) cluster at the University of Birmingham. A dedicated node, equipped with two Intel Xeon Platinum 8570 processors (56 cores each), was employed. Simulations were processed concurrently in batches of 20, with individual runs allocated 4GB of RAM. The duration of each simulation varied with agent count, typically ranging from approximately 45 seconds to 4 minutes.

## 3 Results

### 3.1 Cell invasion and perpendicular

#### fibre realignment increase with chemotaxis bias

In this section, as a baseline study, we investigate the mechanisms by which agent cells reorient the ECM fibres within our model. We systematically varied the ECM sensitivity (*α* in Equation (1)) and chemotaxis bias (*β* in Equation (2)) to analyse their impact on the cell’s capacity to change the fibre orientation over time. Fibre reorientation (S1 text Equation (14)) is directly dependent on the cell’s motility direction, ***d***_*i*,mot_ (Equation (1)), which is, in turn, a function of both ECM sensitivity and chemotaxis bias. We set the reorientation rate to 0.01 min^*−*1^ and the realignment rate to 0.0001 min^*−*1^, as in Metzcar et al. ^13^ . ECM degradation and displacement are deactivated in these simulations so that the dynamics reflected only the effect of chemotaxis bias and ECM sensitivity on migration and remodelling.

We initialised the simulations by randomly placing 100 cells at the bottom of the domain (*y* = *−*500 *µ*m). The ECM was initialised with a density of 0.5 (optimal density for migration, Equation (13) in S1 text), a uniformly random fibre orientation and an initial anisotropy of 0, corresponding to no fibre-fibre alignment correlation (Figure 2A and B, Initial time). As an initial condition, a chemical substrate, representing oxygen, was uniformly distributed throughout the domain, creating an environment of physioxic conditions ^11^ of 38 mmHg (Figure 2C, Initial time). A Dirichlet boundary condition of 38 mmHg was imposed at the top of the domain (*y* = 500 *µ*m) while no-flux boundary conditions were applied on the other sides of the domain. The chemical substrate dynamics were governed by a diffusion coefficient of 100,000 *µ*^2^m min^*−*1^, a decay rate of 0.1 min^*−*1^, and a cell uptake rate of 10 min^*−*1^, with no secretion. These parameters for the chemical microenvironment were used consistently in the subsequent simulation scenarios unless otherwise stated. All the parameters used in the simulations are also listed in the S1 text (Tables 1 to 4). After 96 hours, the cells have invaded the domain, leading to the reorientation of the fibres, an increase in local anisotropy and the establishment of a chemical gradient due to substrate uptake and decay (Figure 2, Final time). In this configuration, oxygen depletion persists near the initial cell region because diffusion from the top boundary competes with decay and cellular uptake, and local consumption can exceed replenishment in dense regions. A video showing the simulations over time can be found in the Supporting information (video S1).

**Figure 2.**
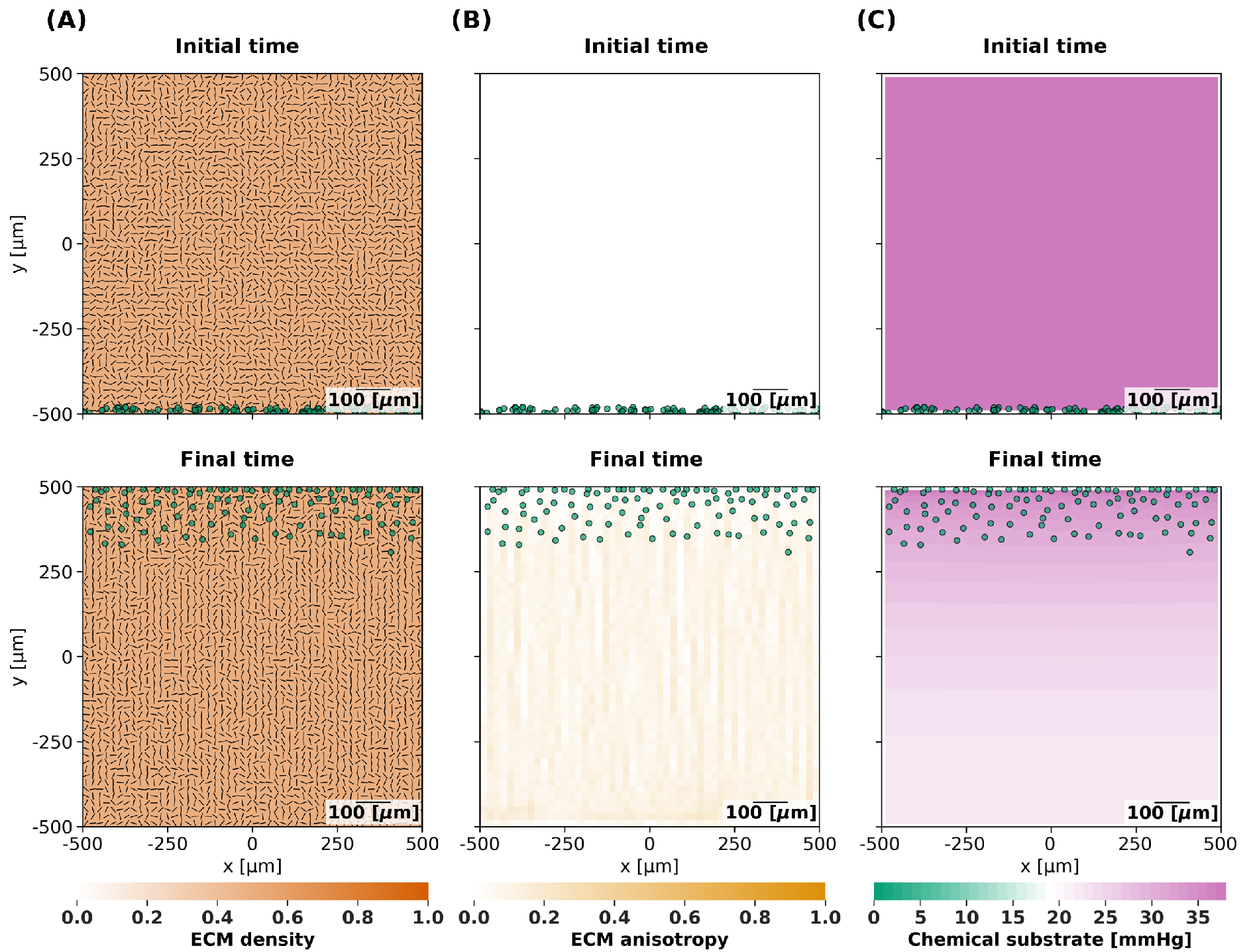
Effect of cell remodelling on ECM and chemical substrate over time on random fibre alignment. Simulation snapshots at initial time (0 h, top row) and final time (96 h, bottom row). Cell agents (green circles) migrate from the bottom to the top of the domain. Panels show (A) fibre orientations (***f***, black segments) with fixed ECM density (*ρ* = 0.5, orange),(B) ECM anisotropy (*a*, yellow), and (C) chemical substrate gradients (***θ***, green to pink).

Figure 3 displays heatmaps illustrating the normalised fibre orientation across the entire domain (Figure 3A) and the total cell invasion (Figure 3B). These metrics are explained in Section 2.5. Figure 3A shows the effects of ECM sensitivity and chemotaxis bias on fibre reorientation. A value of 0.0 indicates that the median of the fibre orientations over the domain is aligned to the *y−*axis, signifying a perpendicular alignment to the tumour boundary. Conversely, a value of 1.0 indicates that the fibres are aligned parallel. Randomly aligned fibres return a value around 0.5. These values provide a reference for understanding the predominant fibre orientation within a given domain. We observe that cells actively realign the fibres perpendicularly when the chemotaxis bias is above 0.4, and the ECM sensitivity is below 0.8. The most significant perpendicular realignment occurs under conditions of maximal chemotaxis bias (*β* = 1.0) and zero ECM sensitivity (*α* = 0.0). In this specific case, the cell’s movement direction, ***d***_*i*,mot_, is solely dictated by the chemotaxis direction, ***d***_chemotaxis_ (Equations (1) and (2)). Consequently, cells exclusively follow the gradient of the chemical substrate originating from the top of the domain, resulting in predominantly movement along the *y−*axis and a subsequent perpendicular reorientation of the ECM fibres. In contrast, for higher ECM sensitivity values, the influence of contact guidance from the fibre orientation ***d***_***f***_ (Equation (4)) becomes more pronounced. Given the initial random fibre orientation and zero initial anisotropy, increased ECM sensitivity leads to less directed cell movement along the *y−*axis, thus maintaining a more random average orientation of the fibres.

**Figure 3.**
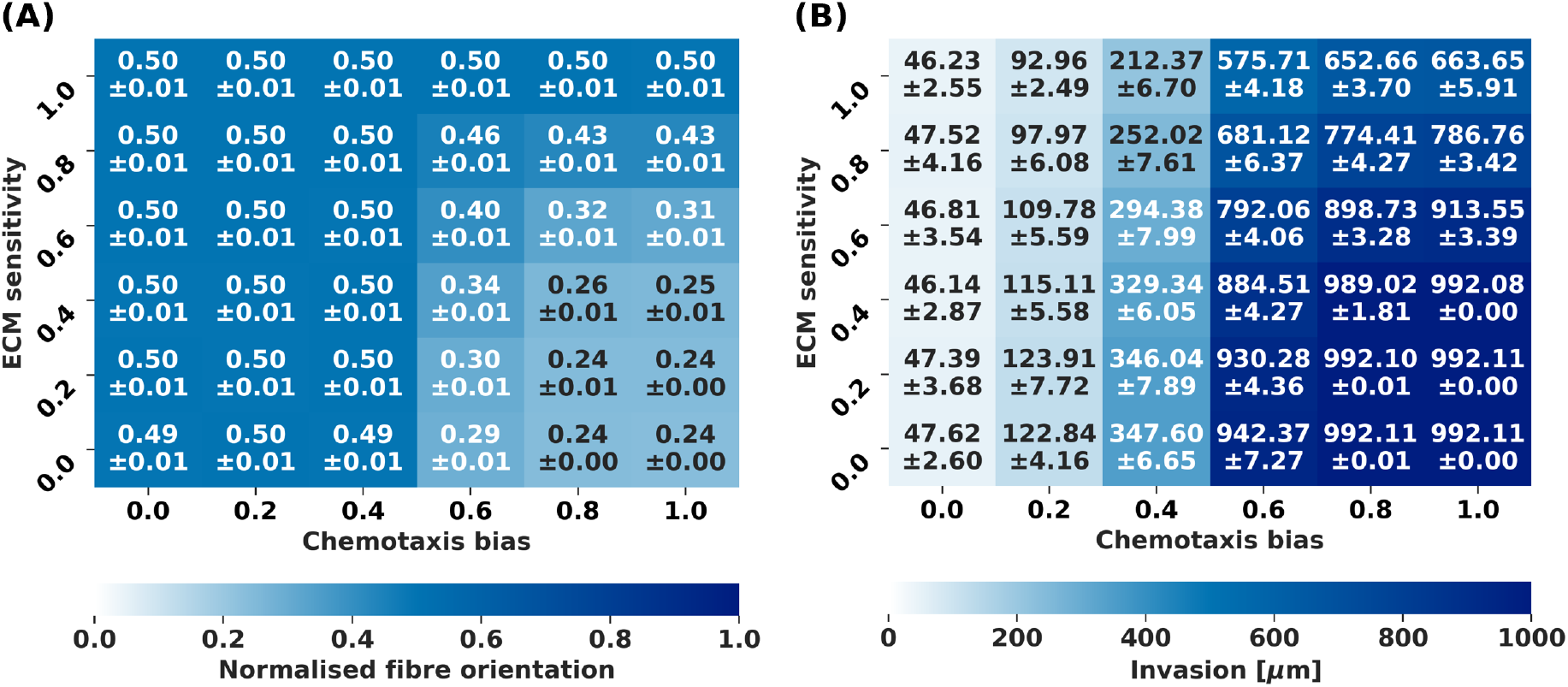
Effect of chemotaxis bias and ECM sensitivity on fibre orientation and invasion (96 h) Heatmaps showing (A) normalised fibre orientation relative to the *y −* axis (90°) and (B) invasion for varying chemotaxis bias (*β*, rows) and ECM sensitivity (*α*, columns) in a domain of size 1000 *×* 1000 *µ*m. Results are averaged over 10 simulations, with mean and standard deviation shown. Metrics are defined in Section 2.5.

Furthermore, Figure 3B demonstrates that cell invasion (explained in Section 2.5) is lower when the chemotaxis bias is below 0.6, as the overall cell movement becomes more random. We also note that, for a constant chemotaxis bias (each column), increasing ECM sensitivity leads to reduced invasion. This effect is attributed to the heightened contact guidance at higher ECM sensitivities, which causes cells to adhere more closely to the initially random fibre orientation, thereby impeding their overall directed migration.

### 3.2 Initial fibre orientation and anisotropy together affect cell invasion

In this section, we explored how the ECM architecture, specifically fibre orientation and anisotropy, influences cancer invasion, drawing inspiration from experimental findings by Provenzano et al. ^20^ . Their work demonstrated that fibres aligned perpendicular to a tumour boundary significantly promote invasion, while parallel alignment suppresses it, highlighting the role of contact guidance in 3D cell migration. Recent computational work in Metzcar et al. ^13^ similarly found that fibres parallel to a tumour surface impair invasion, while fibres perpendicular to the tumour surface facilitate superdiffusive invasion, replicating and extending previous mathematical modelling by Painter ^17^ . We now extend this analysis to more deeply consider the role of the migration parameters with the new cell migration direction implementation (Equations (1), (2), (3) and (4)) by simulating a 2D cross-section. As in the previous simulations, ECM degradation and displacement are deactivated.

We initialised simulations by randomly placing 100 cells within the central region of the domain (*y* = 0 *µ*m), corresponding to the interface between two ECM regions with distinct fibre orientations. The fibres in the top half of the domain were oriented perpendicularly to the tumour boundary (aligned to the *y−*axis), while the fibres in the bottom half were oriented parallel to it (aligned to the *x−*axis). The central region contained randomly oriented fibres (Figure 4A, Initial time). The ECM density was set to 0.5 (Figure 4A, Initial time), and the initial anisotropy was set to 0.5 (Figure 4B, Initial time), introducing a degree of randomness to the fibre orientation (***d***_***f***_, Equation (4)), rather than a perfect alignment, as seen in the experiments^20^. This ensures that cell-fibre interactions include a stochastic component. Finally, a uniform chemical substrate, representing oxygen, was initialised at 38 mmHg across the entire domain (Figure 4C, Initial time), with Dirichlet boundary conditions of 38 mmHg applied at the top (*y* = 500 *µ*m) and bottom of the domain (*y* = *−*500 *µ*m), while the remaining boundaries had no-flux boundary conditions. Other chemical substrate parameters remained consistent with Section 3.1. After 96 hours, the cells have migrated from the origin of the domain (*y* = 0 *µ*m) invading the top and bottom regions of the domain, leading to the reorientation of the fibres, an increase in local anisotropy and the establishment of two chemical gradients, towards the top and bottom of the domain, due to substrate uptake and decay (Figure 4, Final time). With oxygen supplied from the top and bottom boundaries, sustained uptake near invasion fronts can maintain central depletion and limit full re-oxygenation over the simulated interval.

**Figure 4.**
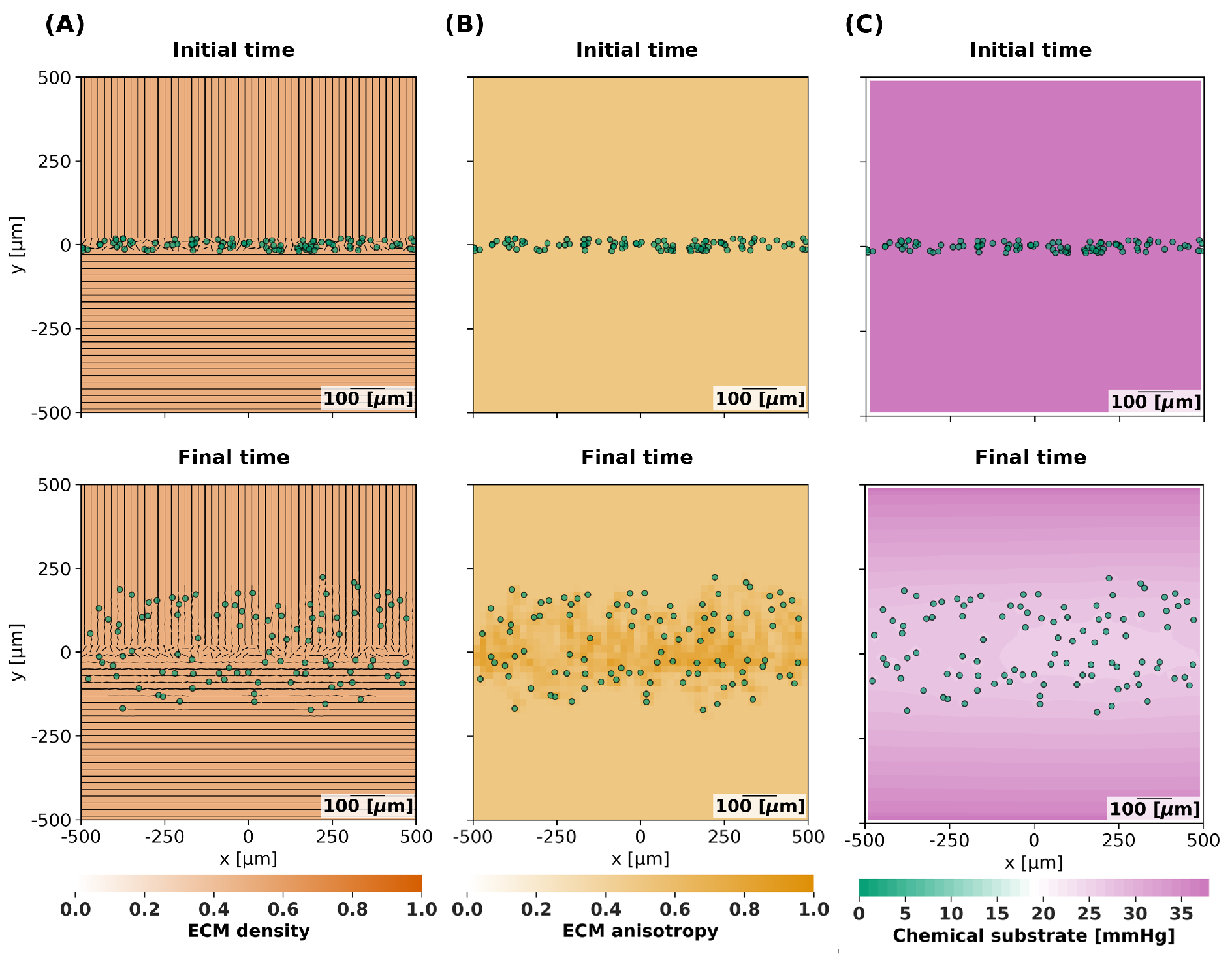
Effect of cell remodelling on ECM and chemical substrate over time on parallel and perpendicular fibre alignments. Simulation snapshots at initial time (0 h, top row) and final time (96 h, bottom row). Cell agents (green circles) migrate from the centre toward the top and bottom of the domain. Panels show (A) fibre orientations (***f***, black segments) with fixed ECM density (*ρ* = 0.5, orange), (B) ECM anisotropy (*a*, yellow), and (C) chemical substrate gradients (***θ***, green to pink).

We investigated the impact of ECM sensitivity (*α* in Equation (1)) and chemotaxis bias (*β* in Equations (2)) on cell invasion. Figure 5 illustrates the spatial distribution of cells (cell count at varying distances from the origin (*y* = 0 *µ*m) of the domain) over the perpendicular and parallel domains after 96 hours. A video showing the simulations over time can be found in the Supporting information (video S2).

**Figure 5.**
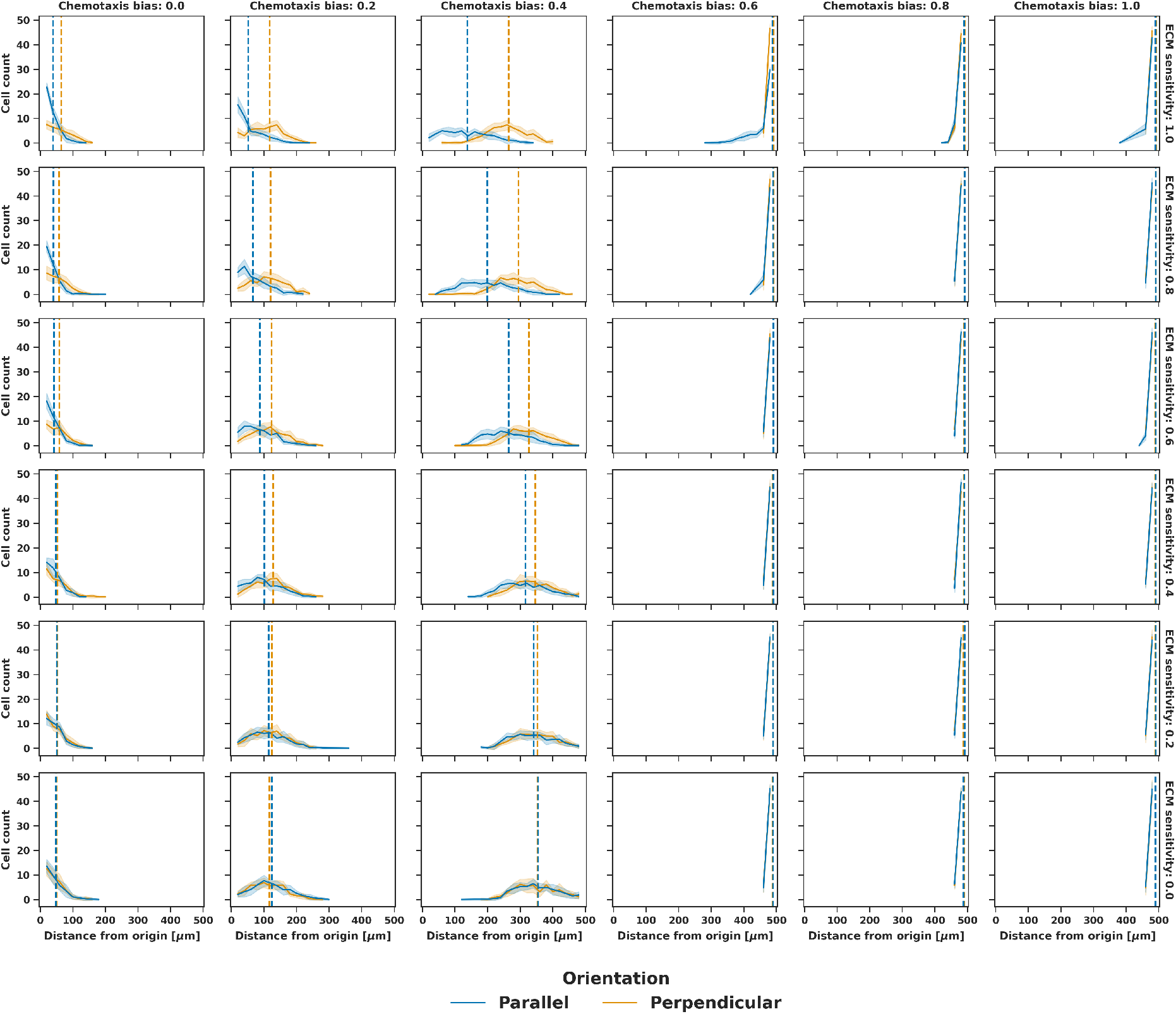
Cell spatial distribution for initial anisotropy 0.5 (96 h) Cell distributions are shown for perpendicular fibre alignment (orange) and parallel fibre alignment (blue). Vertical lines indicate medians. Chemotaxis bias (*β*, columns) and ECM sensitivity (*α*, rows) are varied. Mean (solid line) and standard deviation (shaded area) are averaged over 10 simulations. Metrics are defined in Section 2.5.

When the ECM sensitivity is 0.0, the initial fibre orientation has no noticeable effect on cell migration. That is, the number of cells distributed in the perpendicular (orange) and parallel (blue) domains is equal (Figure 5, bottom row). However, as the ECM sensitivity increases, a significant difference emerges between the two initial fibre alignment conditions at low chemotaxis bias (Figure 5, top rows, chemotaxis bias *β ≤* 0.4). The perpendicular alignment consistently promotes higher invasion compared to the parallel alignment, even in the absence of chemotaxis (*i*.*e*., when chemotaxis bias is 0.0, Figure 5, first column). This highlights the role of physical cues from aligned fibres in guiding or inhibiting invasive behaviour.

Increasing chemotaxis bias enhances invasion for both perpendicular and parallel fibre alignments (Figure 5, left to right). It is worth noting that the chemotaxis bias impacts cell direction even when ECM sensitivity *α* is equal to 1.0 (Figure 5, top row). This is because, while *α* = 1 means cells fully follow the fibre direction (***d***_***f***_, Equation (3)), the sign of this vector is modulated by its dot product with ***d***_pref_ (Equation (4)), which itself depends on chemotaxis (Equation (2)), thereby providing a preferred direction of migration.

Similarly to the case of random fibre orientation explored in Section 3.1, for parallel fibre alignment, a higher ECM sensitivity corresponds to lower overall invasion. Surprisingly, in the perpendicular region for a chemotaxis bias of 0.2 and 0.4 (Figure 5 columns 2 and 3), invasion decreases with increasing ECM sensitivity (Figure 5 orange median line, bottom to top). We would expect that fibres aligned in the same direction of migration facilitate invasion, particularly when contact guidance (ECM sensitivity) is high. This counter-intuitive observation for the perpendicular initialisation is attributed to the inherent randomness in fibre alignment, given by low anisotropy (Equation (4)), which is amplified at higher ECM sensitivities.

To verify this theory, we further investigate the interplay between ECM sensitivity and initial fibre alignment by analysing the effect of varying the initial anisotropy on cell migration, keeping the chemotaxis bias fixed at 0.2. Figure 6 displays the spatial cell distribution for initial anisotropies of 0.3, 0.5 and 0.7, across ECM sensitivities of 0.6, 0.8 and 1.0. A video of the simulations over time can be found in the Supporting information (video S3).

**Figure 6.**
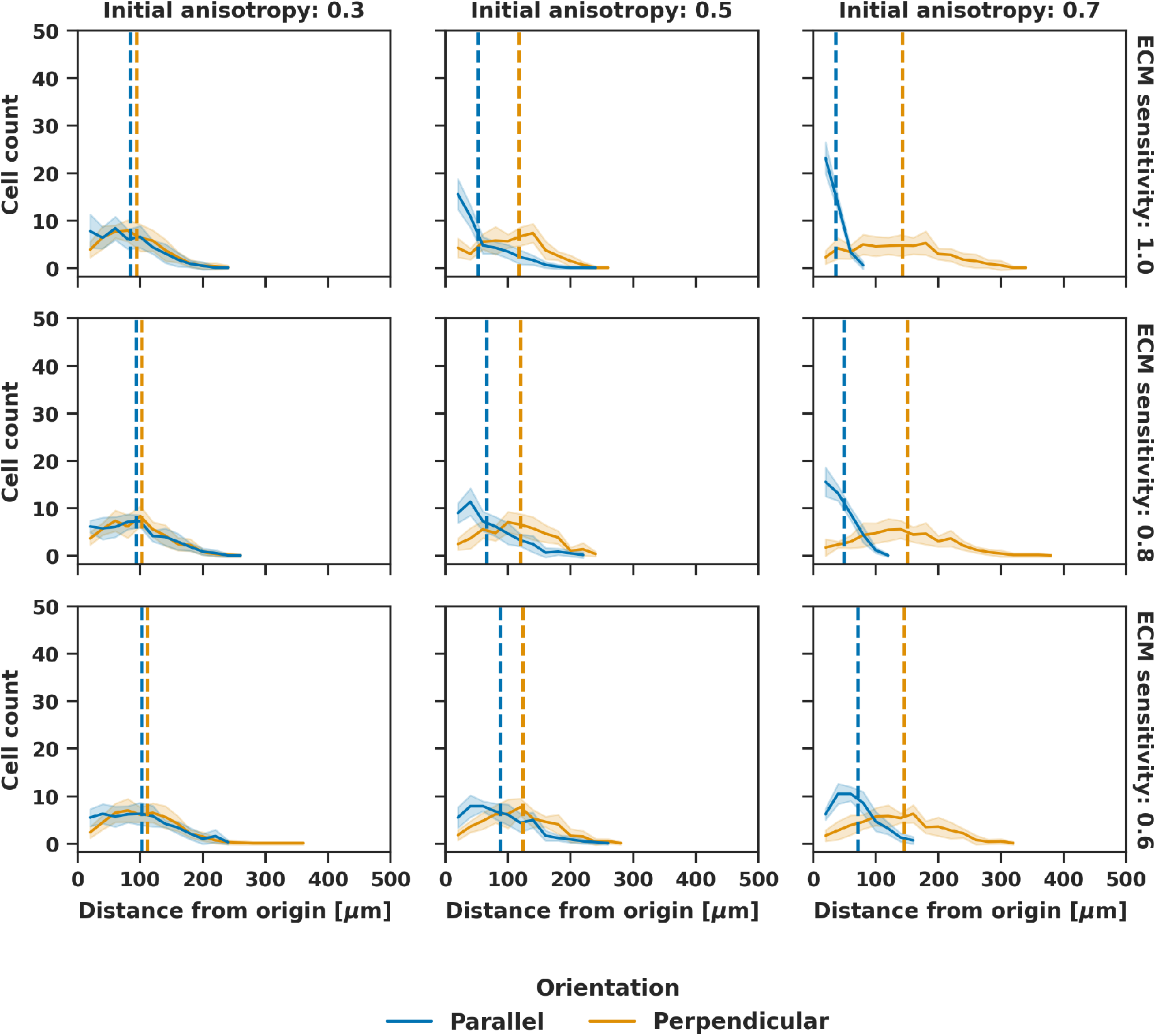
Cell spatial distribution for chemotaxis bias 0.2 (96 h) Cell distributions are shown for perpendicular fibre alignment (orange) and parallel fibre alignment (blue). Vertical lines indicate medians. Initial anisotropy (*a* at *t* = 0, columns) and ECM sensitivity (*α*, rows) are varied. Mean (solid line) and standard deviation (shaded area) are averaged over 10 simulations. Metrics are defined in Section 2.5.

The results underscore the critical role of anisotropy in our model. At an initial anisotropy of 0.3, the initial fibre orientation has a negligible impact on cell migration (Figure 6, first column). This is because an anisotropy close to zero signifies that the fibres within a voxel are largely uncorrelated, essentially mimicking a randomly aligned environment. As initial anisotropy increases, the randomness in the fibre direction vector ***d***_***f***_ (Equation (4)) decreases. High initial anisotropy strengthens the inhibitory effect of parallel fibre alignment and accentuates the differences in cell migration between the perpendicular and parallel initial fibre orientations (Figure 6, left to right). A higher anisotropy leads to more coherent fibre alignment, thereby providing stronger directional cues for cell movement. This leads to decreased invasion in the parallel region and increased invasion in the perpendicular region as the initial anisotropy increases, with the difference becoming more pronounced when the ECM sensitivity is maximal.

These findings emphasise that simply setting an initial fibre orientation is insufficient to influence cell movement. Sufficient fibre-fibre correlation (anisotropy) is essential to translate fibre alignment into directed cell migration within our hybrid model.

Our simulation results resemble the experimental findings by Provenzano et al. ^20^ . They showed that invasion is strongly regulated by ECM architecture: cells invade preferentially into regions where collagen fibres are highly aligned perpendicular to the tumour boundary, whereas invasion is suppressed in regions with low or parallel alignment. In our model, we recover this behaviour under specific parameter regimes in which contact guidance dominates, specifically when ECM sensitivity is high and chemotaxis bias is low, so that perpendicular alignment consistently promotes invasion more than parallel alignment, mirroring the experimental observations. We also reproduce Provenzano et al. ^20^ ‘s contrast between highly aligned and low-alignment matrices, by varying the initial anisotropy: low anisotropy produces behaviour similar to their disordered regions, while high anisotropy reproduces the aligned matrices that facilitate invasion. Based on these findings, we select parameter regimes in which ECM sensitivity, chemotaxis bias, and initial anisotropy reproduce these experimentally observed behaviours for the following simulations.

### 3.3 From compact ECM rims to invasive protrusions: how displacement and degradation reshape spheroid morphology

In this section, we investigate how ECM density remodelling affects the model dynamics. This work introduces a novel mechanism for ECM density displacement (Section 2.3), which, in combination with ECM degradation, can enhance or reduce cell migration and invasion.

We initialised simulations with a cell spheroid of radius 100 *µ*m located at the centre of the domain. Chemotaxis and ECM sensing were both active in this scenario, with a chemotaxis bias of 0.2 and an ECM sensitivity of 0.8 to represent strong contact guidance along ECM fibres. Cell proliferation was enabled with a rate *r*_div_ of 0.00072 min^*−*1^. We set the reorientation rate to 0.01 min^*−*1^ and the realignment rate to 0.0001 min^*−*1^, as in Metzcar et al. ^13^ . The ECM was initialised with a density of 0.7, representing a denser matrix environment that is less favourable for cell movement compared with previous simulations. The fibres are initialised as uniformly random, with a random fibre orientation and an initial anisotropy of 0. The chemical microenvironment was initialised in the same manner as in the previous simulations, with oxygen uniformly distributed at 38 mmHg and Dirichlet boundary conditions applied at all domain boundaries.

We explored displacement and degradation rates spanning several orders of magnitude to capture regimes from negligible to strong remodelling, enabling a systematic characterisation of their combined effects (Figure 7). The parameter sweep revealed that invasion increases as the displacement rate decreases and the degradation rate increases (Figure 7A). At high displacement rates without degradation, spheroids retain a compact morphology, as indicated by low Delaunay mean distance (Figure 7C). Conversely, the largest invasion distances occur when the displacement rate is low, and the degradation rate is moderate to high. Notably, spheroid morphology transitions from compact (low Delaunay mean distance) at high displacement and low degradation, to spread out (high Delaunay mean distance) at low displacement and high degradation. Additionally, spheroid growth is maximal for intermediate displacement rates combined with a high degradation rate (Figure 7B). The biphasic behaviour in the spheroid area was also observed in our previous model, where maximum growth was reached at intermediate degradation rates when the density target value was set to 0^1^. A video showing the simulations over time can be found in the Supporting information (video S4).

**Figure 7.**
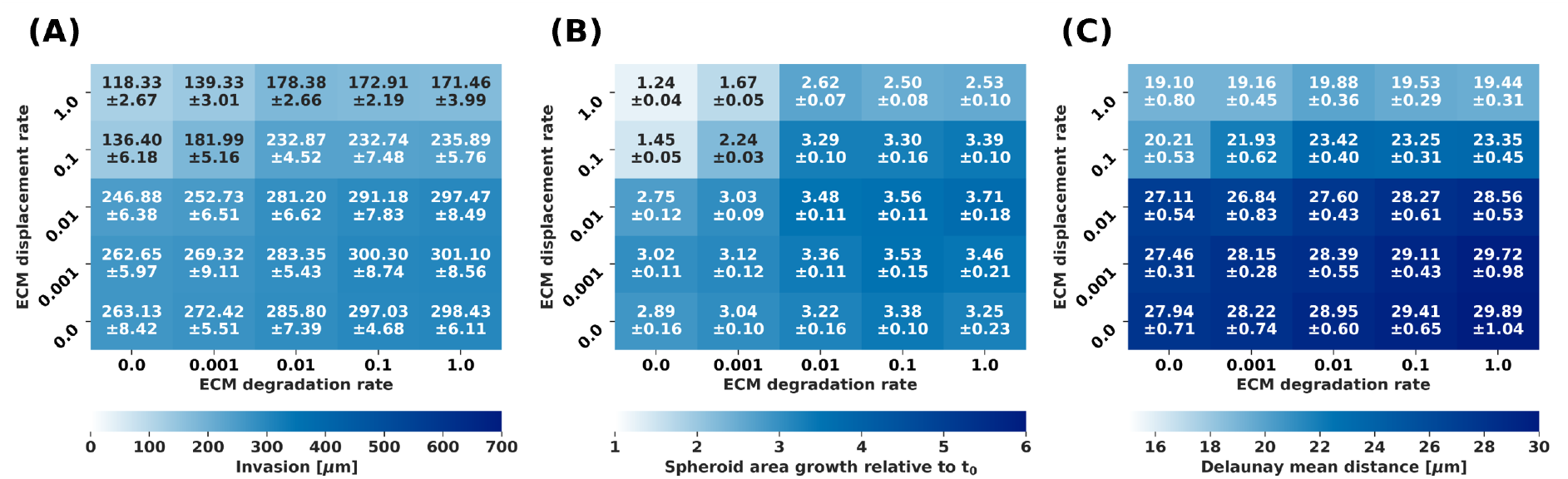
Impact of ECM displacement and degradation on cell migration and proliferation (96 h) Heatmaps show (A) invasion, (B) spheroid growth relative to *t*_0_, and (C) Delaunay mean distance for varying ECM displacement rates (*r*_*d*,disp_, rows) and degradation rates (*r*_*d*,deg_, columns). Results are averaged over 10 simulations, with mean and standard deviation shown. Metrics are defined in Section 2.5.

Simulation snapshots (Figure 8) reveal distinct behaviours across the parameter ranges. We observe the formation of a dense ECM rim around the spheroid at high displacement rates. Because our ECM displacement formulation relocates ECM mass forward during cell migration and expansion-driven pushing, repeated cellular interactions lead to local matrix accumulation. If degradation is insufficient to clear this barrier, the rim restricts further outward invasion. The phenomenon of ECM rim formation has been observed experimentally Lee et al. ^9^ . High degradation rates mitigate this effect by removing the accumulated matrix, allowing for deeper invasion. While moderate degradation promotes the formation of multicellular protrusions and defined invasion tracks, very high degradation leads to broad cell dissemination and maximal expansion. Thus, our model demonstrates how the crosstalk between ECM displacement and degradation dictates the invasive behaviour of cells. Full simulation grids showing ECM density, anisotropy, and chemical substrate distributions without cells are provided in the Supporting information (Figures S1–S3).

**Figure 8.**
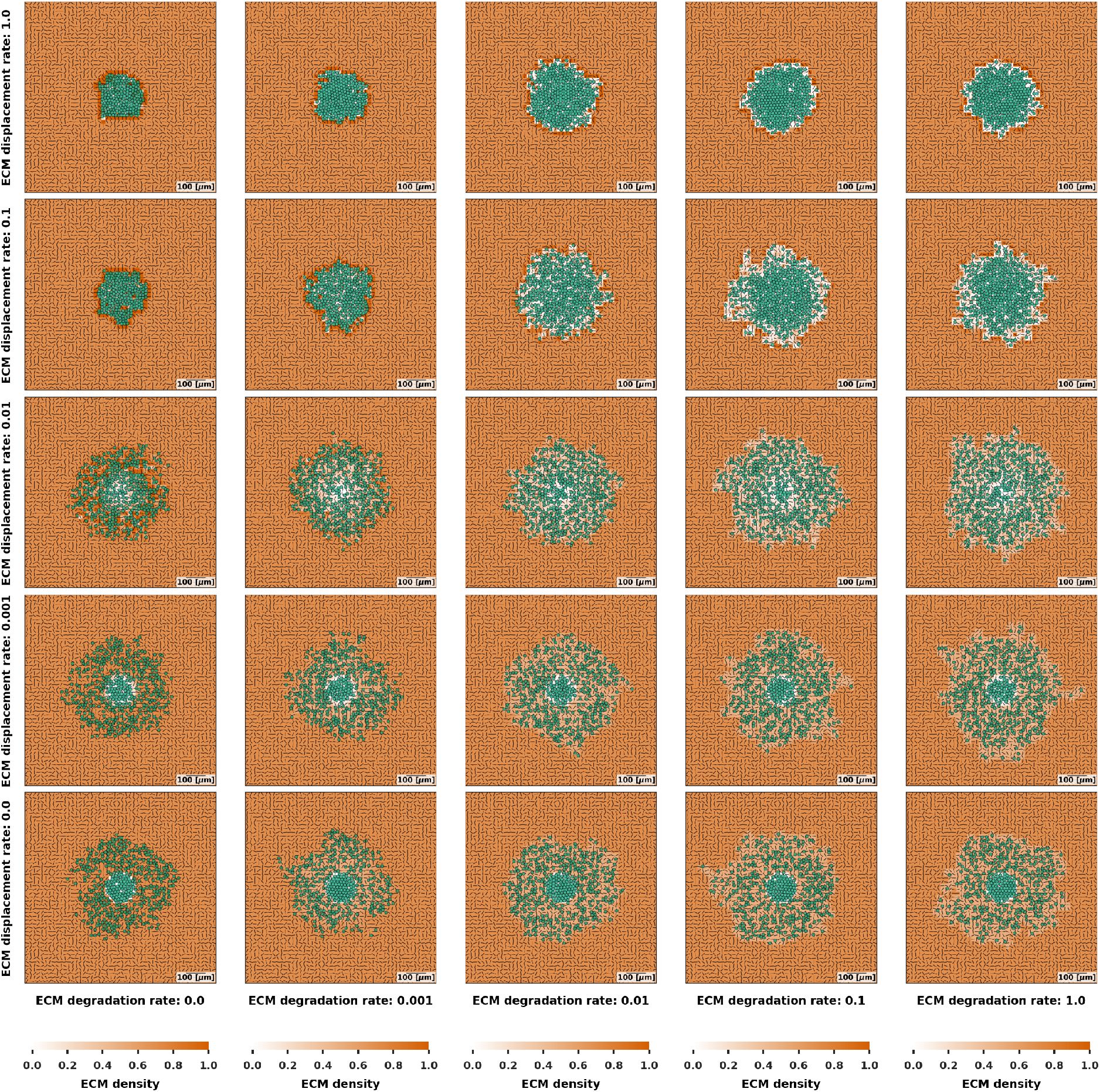
Simulation visualisations for spheroid growth and invasion (96 h) Figures show cancer cells (green), fibre orientations (black segments) and ECM density (orange) for varying ECM displacement rates (*r*_*d*,disp_, rows) and degradation rates (*r*_*d*,deg_, columns).

### 3.4 Impact of fibre reorientation rate and initial fibre architecture on spheroid invasion and growth

Here, we investigated the interplay between ECM reorientation rates and initial fibre orientations on cancer spheroid migration and proliferation. This is based on experimental observations by Su et al. ^30^, which highlighted the influence of initial fibre orientation on spheroid invasion.

We initialise the simulations as in Section 3.3 and we set the degradation rate to *r*_*d*,deg_ = 0.01 min^*−*1^ and the displacement rate to *r*_*d*,disp_ = 0.1 min^*−*1^. We explored three distinct initial fibre orientations: tangential, random, and radial. We set the initial anisotropy to 0.7 to have some randomness in the fibre direction vector (***d***_***f***_, Equation (4)) that resembles the realistic fibre structures.

Figure 9 summarises the effects of varying reorientation rates on invasion, Delaunay mean distance, and spheroid growth relative to initial time (explained in Section 2.5) for the three initial fibre orientations (tangential, random and radial). The colorbar scale for invasion in Figure 9A extends to 700 *µ*m, which represents the maximum possible invasion distance from the origin (centre) of the domain given the 1000*×*1000 *µ*m domain size. A video of the simulations over time can be found in the Supporting information (video S5). Full simulation grids showing ECM density, anisotropy, and chemical substrate distributions without cells are provided in the Supporting information (Figures S4–S6).

**Figure 9.**
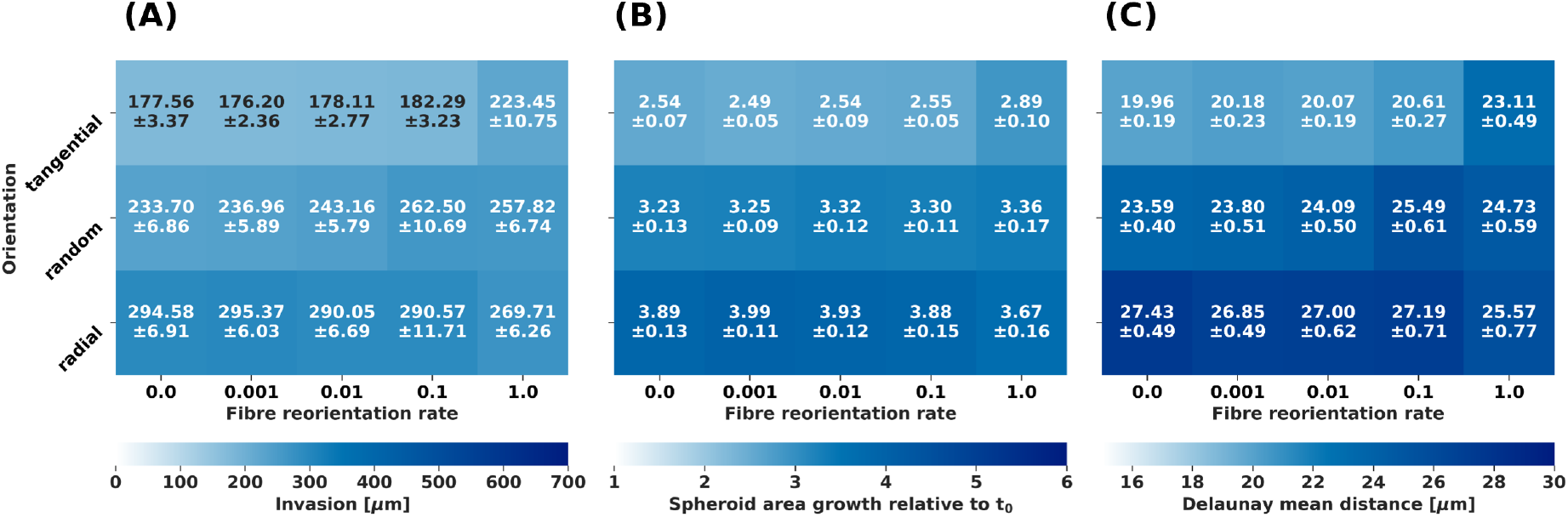
Impact of fibre reorientation rate and initial fibre orientation on cell migration and proliferation (96 h) Heatmaps show (A) invasion, (B) spheroid growth relative to *t*_0_, and (C) Delaunay mean distance for tangential, random, and radial initial fibre orientations (rows) at varying fibre reorientation rates (*r*_*f*0_, columns). Results are averaged over 10 simulations, with mean and standard deviation shown. Metrics are defined in Section 2.5.

At lower reorientation rates, a clear dependency on initial fibre orientation emerges. Compared to the random initialisation, invasion and proliferation are significantly inhibited for the tangential initialisation, while they are enhanced for radial alignment (Figure 9A and B). This trend can be visually observed in Figure 10. Spheroids with tangential initial fibre orientation (Figure 10, top row) maintain a more rounded morphology with minimal or no protrusions, indicating restricted outward migration. Conversely, radial alignment (Figure 10, bottom row) actively promotes collective invasion and the formation of distinct finger-like protrusions, which are extensions from the central spheroid that enable cells to explore and invade the surrounding ECM more extensively, facilitating their invasion into the matrix.

**Figure 10.**
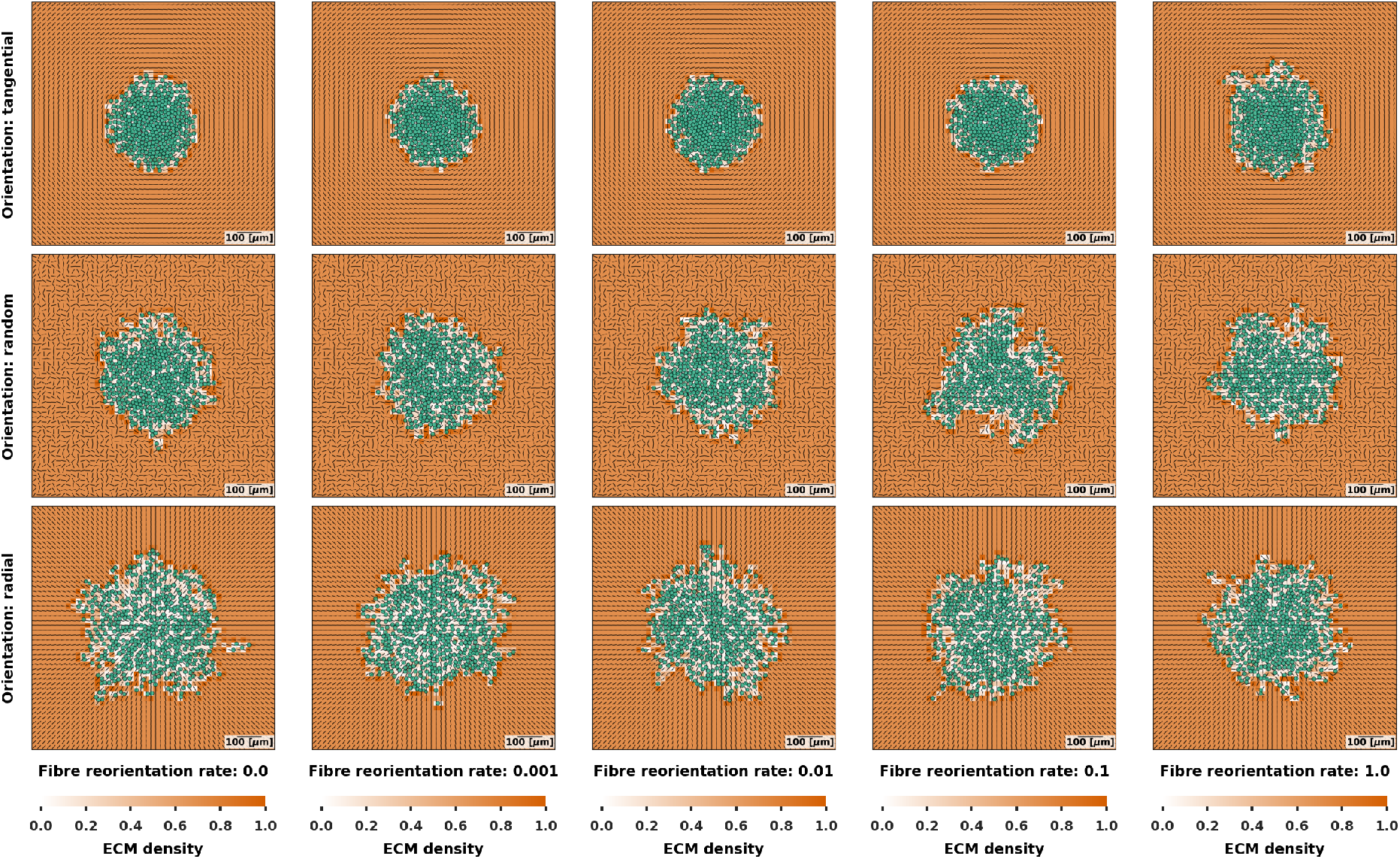
Simulation visualisations for spheroid growth and invasion (96 h) Figures show cancer cells (green), fibre orientations (***f***, black segments) and ECM density (*ρ*, orange) for different reorientation rates (*r*_*f*0_) and initial fibre alignments: tangential (top row), random (central row), and radial (bottom row).

However, a critical shift occurs as the reorientation rate increases (*i*.*e*., reorientation rate greater than 0.1). For the tangential fibre orientation, we observe a notable increase in invasion (Figure 9A, top row) and a moderate increase in growth (Figure 9B, top row) by an increase in *r*_*f*0_. This occurs because a higher reorientation rate allows the cells to rapidly modify the fibre alignment. When fibres are initially tangential, they inhibit migration, due to high contact guidance (*α* = 0.8). Thus, an increased capacity for fibre reorientation (Supporting information, Figure S4, top row) allows the cells to actively modify these impeding fibres, enabling more efficient migration and invasion. Conversely, when fibres are initially radially aligned, increasing the reorientation rate leads to a decrease in invasion (Figure 9A, bottom row) and a slight reduction in growth (Figure 9B, bottom row). In this optimal radial configuration, any deviation from the ideal radial alignment (here caused by reorientation, Supporting information, Figure S4, bottom row) effectively reduces invasion. Interestingly, when the initial fibre orientation is random, invasion and growth show a mild increasing trend with the fibre reorientation rate, as seen in Figure 9, central row. This suggests that for an initially random ECM, the induced changes in fibre orientation due to cellular reorientation are insufficient to significantly impact overall cell invasion or growth for chemotaxis bias 0.2 (Supporting information, Figure S4, central row). The observed trends, particularly a partial convergence of outcomes across different initial orientations at high reorientation rates (Figures 9 and 10, last columns), can be further understood by considering the principles discussed in Section 3.1 regarding the interplay of ECM sensitivity, chemotaxis bias, and fibre reorientation.

The Delaunay mean distance, which serves as a measure of cell packing density, generally reflects the invasion and growth patterns (Figure 9C). Higher invasion and growth often correlate with an increased Delaunay mean distance, indicating a less dense and more spread-out cellular configuration as cells migrate and proliferate into the ECM. Conversely, reduced invasion and growth, particularly with tangential initial fibres at low reorientation rates, lead to a smaller Delaunay mean distance, suggesting tighter cell clustering.

Our simulation results align qualitatively with experimental observations of cancer spheroid invasion in various ECM environments. For instance, the inhibited invasion observed with tangential fibre orientations at low reorientation rates resembles the constrained growth of certain cancer cell lines (*e*.*g*., MCF7) when embedded in matrices with circumferential fibres, as reported by Su et al. ^30^ . The enhanced invasion in radial environments and with increased reorientation capacity for initially restrictive architectures is consistent with the ability of highly invasive cancer cells (*e*.*g*., T47D) to remodel their microenvironment and form invasive protrusions or finger-like projections, even in initially challenging fibre arrangements^30^. This suggests that the capacity for ECM remodelling through reorientation is a key determinant of invasive potential, particularly when overcoming intrinsically inhibitory fibre architectures. Our computational experiments appear to capture these phenomena, providing valuable insights into the biophysical mechanisms underlying cancer cell invasion in relation to ECM fibre architecture.

## 4 Discussions

In this study, we presented a hybrid computational model to investigate the effects of fibre architecture and ECM remodelling on cancer invasion and proliferation. Our results show that the ECM architecture can play a fundamental role in regulating invasion, and that contact guidance and chemotaxis can synergistically direct cell migration. These model-driven insights can drive future experiments for validation, while also predicting specific tumour invasive behaviours that may be observed in experimental and clinical studies.

Consistent with experimental observations by Provenzano et al. ^20^ and Su et al. ^30^, our simulations predict that initial fibre orientation significantly affects invasion outcomes. For high ECM sensitivity and low chemotaxis bias, perpendicularly (Figure 5, orange line) or radially (Figure 9, bottom row) aligned fibres promoted invasion, while parallel (Figure 5, blue line) or tangential (Figure 9, top row) alignment inhibited it. Furthermore, within our model, as a consequence of how we implemented anisotropy, we found that without sufficient anisotropy, even a favourable fibre alignment failed to significantly influence cell trajectories, suggesting that effective directional guidance requires both orientation and structural coherence (high initial anisotropy). This aligns with reported experimental findings by Provenzano et al. ^20^ and Su et al. ^30^, where different techniques were employed to experimentally align collagen fibres, demonstrating that distinct fibre architectures (*e*.*g*., highly aligned versus disorganised) directly influenced cancer cell migration and invasion.

Our model also shows that while ECM sensitivity can enhance contact guidance, it can reduce overall invasion, even when the fibre orientation is aligned with the direction of invasion of the cells. This is attributable to the stochasticity in fibre directions (Equation (3)), which introduces noise into the contact guidance signal, particularly at low anisotropy. As a result, excessive reliance on ECM cues without sufficient fibre alignment can restrict effective migration, especially when chemotactic cues are weak. Specifically, we observed that high ECM sensitivity amplifies the inhibitory effect of parallel fibre alignment, particularly at high initial anisotropy, but does not increase invasion when fibres are perpendicular to the tumour boundary. In these cases, cells still require directional cues to determine a preferred migration direction, making chemical gradients essential for guiding migration. This interplay between chemical and mechanical cues suggests that tumours may switch between chemotaxis-dominated and contact guidance-dominated invasion modes depending on local ECM organisation, a hypothesis that could be tested experimentally by modulating collagen alignment while applying controlled chemotactic gradients.

Our results also highlight the distinct roles of ECM degradation and density displacement in regulating invasion. In simulations where degradation dominated, cells progressively cleared the matrix and generated migration tracks that facilitated further and sparse invasion. In contrast, density displacement redistributed matrix material without removing it, often leading to the formation of dense rims surrounding the tumour boundary, impeding cell migration and leading to a more compact spheroid morphology. The balance between these mechanisms, therefore, plays an important role in shaping invasion dynamics and may help explain experimentally observed accumulation of collagen around expanding tumour spheroids ^9^.

Our results also exhibit conceptual similarities to flocking and active-matter models of collective migration ^33^. In both settings, simple local interaction rules can produce emergent coordinated behaviours such as aligned motion, leader-follower organisation, and the formation of protrusive invasion structures. However, in classical flocking models, alignment typically arises through direct neighbour-neighbour interactions between agents. In contrast, alignment in our framework is mediated indirectly through a remodelled environment: cells align with ECM fibres rather than directly with neighbouring cells. This environmental feedback loop, where cells follow fibres while simultaneously reorienting and redistributing them, places our model at the interface between flocking dynamics and mechanically remodelled active media. As such, analytical tools from flocking theory may provide a useful framework for analysing the stability, persistence, and branching of invasive protrusions observed in our simulations.

Our findings also highlight several model limitations and opportunities for future development. The current model assumes monotonic increases in anisotropy, which limit its ability to capture reversible or disorganising ECM remodelling observed in real tumours. This is particularly relevant given that high anisotropy in our model slows and ultimately blocks reorientation of the ECM (Equation (14). Introducing mechanisms for anisotropy reduction could improve realism and enable the simulation of ECM de-alignment.

In addition, fibre orientation and anisotropy updates are currently localised to the voxel closest to a cell’s front, ignoring the spatial continuity of fibres across neighbouring regions. Future extensions could implement a spatially correlated remodelling scheme to reflect the interconnected nature of ECM networks better and facilitate more realistic ECM remodelling and invasion dynamics. Such extensions would also allow local remodelling to propagate through the matrix and generate long-range alignment patterns, as observed experimentally in tumour-associated collagen signatures.

Matrix remodelling in the current model is also permanent: once the matrix is relocated or the fibres reoriented, no elastic force restores it to a reference configuration. Thus, ECM remodelling represents plastic remodelling rather than elastic deformation. Furthermore, displacement currently affects only ECM density and does not influence fibre orientation and anisotropy. Experimental studies have shown that mechanical forces generated during spheroid expansion not only produce an accumulation of ECM, forming the characteristic dense rim surrounding the spheroid, but also alter fibre orientation. Compressive forces near the spheroid boundary tend to generate tangential fibre alignment, whereas traction forces at longer distances promote radial alignment. Coupling matrix displacement to fibre reorientation would therefore provide a natural extension of the model and enable pushing-induced compaction near the spheroid edge to generate circumferential alignment while traction forces promote radial alignment, more closely reproducing experimentally observed ECM architectures.

ECM density, fibre orientation, and anisotropy are also closely interconnected. Changes in density can alter the ability of cells to reorient fibres, an effect that is further modulated by the degree of fibre cross-linking. Cross-linking was not explicitly included in the present model. In biological tissues, higher density and cross-linking typically increase matrix stiffness and resistance to reorientation, while also modifying how forces propagate through the fibre network. Incorporating density- and cross-linking-dependent reorientation rates, or allowing anisotropy to relax in regions of low cross-linking, could enable the model to capture a broader spectrum of ECM behaviours, including strain-induced alignment and partial structural recovery following mechanical unloading.

The current model also does not include matrix deposition or cell death. As a result, ECM can only be removed or displaced but not replenished, and all cells remain viable throughout the simulations. Incorporating matrix deposition would allow tumour or stromal cells to reinforce migration tracks or rebuild obstructive barriers within the matrix. Similarly, including cell death and necrosis could alter local mechanical pressures and oxygen consumption, potentially influencing both invasion dynamics and ECM remodelling over longer timescales.

Finally, the model is also subject to technical and dimensional constraints. The ECM voxel size is currently set to 20 *µ*m, which is slightly larger than the maximum diameter of the cell agents. Smaller voxels would provide higher resolution interactions. However, reducing voxel size increases the computational cost. Our simulations are in a constrained 3D domain, representing a 2D cross-section, despite the inherently 3D nature of tumour-ECM interactions. While our model conceptually addresses 3D effects on cell migration, all simulations were conducted on 2D slices. Additionally, the underlying ECM framework ^13^ supports only constrained 3D simulations. This simplification restricts the ability to capture unconstrained 3D cell movement and ECM effects. Furthermore, cell volume is approximated and cell shapes are not explicitly modelled, which may impact the accuracy of predicted cell-ECM interactions and migration patterns. Similarly, subcellular mechanisms of cell motility, such as protrusion formation (*e*.*g*. lamellipodia or filopodia) and adhesion turnover with the extracellular matrix, are not explicitly represented. Instead, these processes are coarse-grained into the effective motility velocity used in the model. The insights gained from this pseudo-2D model will guide future development of fully 3D simulations with more realistic cell morphology. Nevertheless, we expect that a constrained 3D representation would still capture the essential dynamics of spheroid growth and collective invasion. Modelling ECM fibre architecture in fully unconstrained 3D environments presents substantial technical and experimental challenges, particularly when attempting to compare simulations with imaging data. In practice, most experimental analyses of collagen organisation rely on two-dimensional slices or projections, making direct validation of fully 3D fibre networks difficult. For this reason, intermediate modelling approaches that incorporate constrained 3D geometries while preserving experimentally measurable fibre features may provide a practical path toward quantitative validation.

In summary, our hybrid model offers a powerful framework for investigating the complex and dynamic interactions between cancer cells and the ECM. It provides mechanistic insights into how ECM structure could influence invasion, and how cellular remodelling could reshape this structure in return. By closely aligning with experimental findings and highlighting new hypotheses regarding the interplay between chemotactic guidance, fibre anisotropy, and mechanically induced ECM remodelling, this work lays a strong foundation for future integrative studies in computational oncology. In particular, elucidating how key invasive behaviours can be modulated through purely mechanical means that are often overlooked in systems biology approaches that focus more narrowly on responses to diffusible ligands. As experimental and mathematical biology converge and approach consensus on describing tumour-ECM contact interactions, the mathematical framings in this paper could be integrated into emerging multicellular standards ^28;8^.

## Author contributions

MB Conceptualization, Formal analysis, Methodology, Project administration, Software, Visualization, Writing - original draft, review & editing JM: Supervision, Writing - review & editing PM: Writing - review & editing PK: Conceptualization, Project administration, Supervision, Writing - review & editing FS: Conceptualization, Project administration, Supervision, Writing - review & editing

## Acknowledgments

We acknowledge the use of Grammarly (version 14.1248.0) and ChatGPT-5 for text editing and to identify improvements in writing style. The computations described in this paper were performed using the University of Birmingham’s BlueBEAR HPC service, which provides a High Performance Computing service to the University’s research community. See http://www.birmingham.ac.uk/bear for more details.

## Data availability statement

The datasets presented in this study are available in online repositories. The names of the repository/repositories and accession number(s) can be found below: https://github.com/Margherita-Botticelli/PhysiCell-cancer-ecm-fibers.

