## Supplementary material for "The role of contact guidance and ECM remodelling in cancer invasion: a computational study": S1_text

#### 1 Supplementary methods

As in our previous work in Botticelli et al.[2], we employ a pseudo-2D (or constrained 3D) model representing a $z$-slice image, akin to the central cross-section of *in vitro* experimental spheroid data (Figure 1A). The cells are modelled as spherical agents with a maximum volume $V$ and corresponding radius $R$, and interact with a single layer of ECM voxels. While cell agents can move freely in the $x$- and $y$-directions within this layer, their movement is constrained in the $z$-direction, effectively creating a constrained 3D model. This framework was chosen to facilitate direct comparison between simulated and experimental data, as experimental images are often captured from $z$-slices in the case of cancer spheroids, particularly the central $z$-slice.

We reutilised the voxel selection method developed in Botticelli et al.[2] for cell movement and ECM remodelling. Invasive cancer cells form outward protrusions (*e.g.*, filopodia and invadopodia), crucial for mechanically and chemically remodelling ECM fibres, often secreting matrix metalloproteinases (MMPs)[7;3]. To reflect this biology, cells interact with the ECM voxel located at their "leading edge" rather than the voxel containing their centre ($\boldsymbol{x}_i$), in contrast to the approach in Metzcar et al.[11]. For a cell at position $\boldsymbol{x}_i$ with direction of motion $\boldsymbol{d}_i$, we define a point on the cell surface in the direction of motion as $\boldsymbol{p}_i = \boldsymbol{x}_i + R_i\boldsymbol{d}_i$, where $R_i$ is the cell's mean radius (Figure 1B). The voxel at the "cell front" is the nearest voxel to the point $\boldsymbol{p}_i$. When the cell is moving (*i.e.*, $\boldsymbol{v}_i \neq \boldsymbol{0}$), all functions that involve the ECM (ECM remodelling in Section 1.3 and cell motility in Section 1.2.2 and Section 2.2 of the main text), are computed at the "cell front", with the only exception of ECM density displacement (in the main text Section 2.3, see Figure 1). If a cell is stationary ($\boldsymbol{v}_i = \boldsymbol{0}$), we instead use the nearest voxel to the cell's position $\boldsymbol{x}_i$.

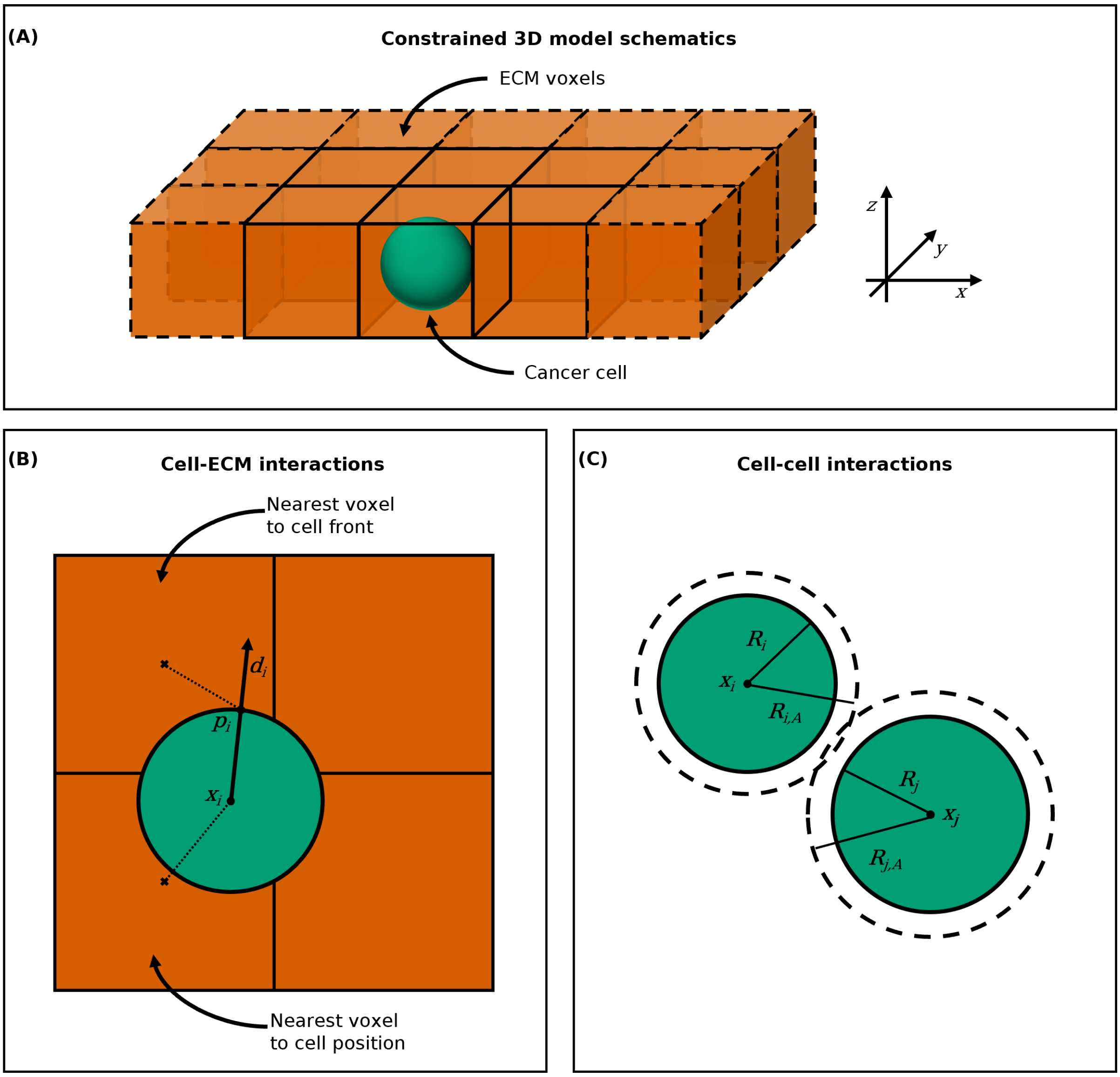


Figure 1: Figure from Botticelli et al.[2]. (A) Constrained 3D model schematics. The ECM voxels are represented as orange cubes and the cancer cell as a green sphere. The cancer cell agent moves freely within the ECM voxels in the $x-$ and $y$-directions, but not in the $z$-direction. (B) Cell-ECM interactions computation. The cell (green) interacts with the ECM voxel (orange) whose centre $(\times)$ is nearest to either its position $\boldsymbol{x_i}$ or its front $\boldsymbol{p}_i$, determined as the point on the cell surface in its direction of motion $\boldsymbol{d}_i$ (C) Cell-cell interactions computation. Cell $C_i$ on the left has a radius $R_i$ and an interaction radius $R_{i,A}$ and cell $C_j$ on the right has a radius $R_j$ and an interaction radius $R_{j,A}$. The two cells interact when their distance is less than $R_{i,A} + R_{j,A}$.

#### 1.1 Chemical microenvironment

The chemical microenvironment consists of diffusible chemical substrates with corresponding diffusion coefficients, decay rates, initial conditions and boundary conditions. PhysiCell uses BioFVM[5], a fast multi-substrate diffusion solver, to simulate chemical microenvironment dynamics using reaction-diffusion-decay partial differential equations (PDEs). It relies on a Cartesian mesh and uses the finite volume method. In addition to diffusion and decay, substrates can be secreted and taken up by the cell agents. The overall PDE, which describes the dynamics of the substrates, is as follows:

$$\frac{\partial \boldsymbol{\theta}}{\partial t} = \underbrace{\boldsymbol{D}\nabla^2\boldsymbol{\theta}}_{\text{diffusion}} - \underbrace{\boldsymbol{\lambda}\boldsymbol{\theta}}_{\text{decay}} + \underbrace{\boldsymbol{S}(\boldsymbol{\theta}^* - \boldsymbol{\theta})}_{\text{bulk source}} - \underbrace{\boldsymbol{U}\boldsymbol{\theta}}_{\text{bulk uptake}} + \underbrace{\sum_i \delta(\boldsymbol{x} - \boldsymbol{x}_i) V_i \left[ \bar{\boldsymbol{S}}_i (\bar{\boldsymbol{\theta}}_i^* - \boldsymbol{\theta}) - \bar{\boldsymbol{U}}_i \boldsymbol{\theta} \right]}_{\text{secretion and uptake by cells}}. \quad (1)$$

Variables and parameters presented in Equation (1) are described in Table 1.

| Symbol | Meaning | Dimensions |
|---|---|---|
| $\boldsymbol{\theta}$ | vector of substrate densities (or concentrations) | substance/length$^3$ |
| $\boldsymbol{D}$ | vector of diffusion coefficients | length$^2$/time |
| $\boldsymbol{\lambda}$ | vector of decay rates | 1/time |
| $\boldsymbol{S}$ | vector of secretion rates | 1/time |
| $\boldsymbol{\theta}^*$ | vector of secretion saturations | substance/length$^3$ |
| $\boldsymbol{U}$ | vector of uptake rates | 1/time |
| $V_i$ | volume of cell $C_i$ | length$^3$ |
| $\boldsymbol{x}_i$ | cell $C_i$'s position (centre) | length |
| $\bar{\boldsymbol{S}}_i$ | vector of cell $C_i$'s secretion rates | 1/time |
| $\bar{\boldsymbol{\theta}}_i^*$ | vector of cell $C_i$'s secretion saturations | substance/length$^3$ |
| $\bar{\boldsymbol{U}}_i$ | vector of cell $C_i$'s uptake rates | 1/time |

Table 1: Description of the variables and parameters in the reaction-diffusion-decay PDE in Equation (1)

#### 1.2 Cell motion

Cell motion in PhysiCell[6] is modelled with the equation:

$$m_i \dot{\boldsymbol{v}}_i = \sum_{C_j \in \mathcal{N}_i} (\boldsymbol{F}_{cca}^{ij} + \boldsymbol{F}_{ccr}^{ij}) + \boldsymbol{F}_{\text{mot}}^i + \boldsymbol{F}_{cmr}^i + \boldsymbol{F}_{drag}^i. \quad (2)$$

Here, $m_i$ represents the mass of a cell $C_i$, $\boldsymbol{v}_i$ is its total velocity, $\boldsymbol{F}_{cca}^{ij}$ and $\boldsymbol{F}_{ccr}^{ij}$ are the cell-cell adhesion and repulsion forces between the cell $C_i$ and the cells $C_j$ in its neighbouring set $\mathcal{N}_i$, given by the cells within interaction distance of each other, respectively, $\boldsymbol{F}_{\text{mot}}^i$ is the motility force, $\boldsymbol{F}_{cmr}^i$ is the cell-ECM repulsion (or resistance) force, and $\boldsymbol{F}_{drag}^i$ is a dissipative, drag-like force. The drag force is modelled as:

$$\boldsymbol{F}_{drag}^i = -\nu \boldsymbol{v}_i, \quad (3)$$

where $\nu$ is the drag coefficient. Applying the inertialess assumption as in Drasdo et al.[4] ($m_i \dot{\boldsymbol{v}}_i \approx \mathbf{0}$ ), we can solve the equation for $\boldsymbol{v}_i$ explicitly

$$\boldsymbol{v}_i = \frac{1}{\nu} \left( \sum_{j \in \mathcal{N}_i} (\boldsymbol{F}^{ij}_{cca} + \boldsymbol{F}^{ij}_{ccr}) + \boldsymbol{F}^{i}_{\mathrm{mot}} + \boldsymbol{F}^{i}_{cmr} \right). \tag{4}$$

Under the inertialess assumption, each force term can be interpreted as specifying the cell's terminal velocity (given drag), assuming all other forces are zero. Therefore, we can write the total velocity $\boldsymbol{v}_i$ of a cell $C_i$ as

$$\boldsymbol{v}_i = \boldsymbol{v}_{i,cca} + \boldsymbol{v}_{i,ccr} + \boldsymbol{v}_{i,\mathrm{mot}} + \boldsymbol{v}_{i,cmr}, \tag{5}$$

where $\boldsymbol{v}_{i,cca}$ and $\boldsymbol{v}_{i,ccr}$ are the cell-cell adhesion and repulsion velocities as a result of cell-cell adhesion and repulsion forces, and $\boldsymbol{v}_{i,\mathrm{mot}}$ and $\boldsymbol{v}_{i,cmr}$ are the motility and repulsion velocities as a result of motility and repulsion forces. We note that, unlike prior work by Ghaffarizadeh et al.[6] and Metzcar et al.[11], cell migration is not modelled as an imposed additional force term, but rather as the net balance of cell-ECM force interactions.

##### 1.2.1 Cell-cell adhesion and repulsion

As in Botticelli et al.[2], we use the built-in functions in PhysiCell for cell-cell adhesion and repulsion[9;6;11]. Let $R_i$ and $R_j$ be the equivalent radii of the cells $C_i$ and $C_j$ respectively, and let $R_{i,A}$ and $R_{j,A}$ be the interaction radii. When the distance between two cell centres $|\boldsymbol{x}_i - \boldsymbol{x}_j|$ is less than their interaction distance $R_{i,A} + R_{j,A}$, we consider cell-cell adhesion to be activated, and the cell agents start pulling each other

$$\boldsymbol{F}^{ij}_{cca} = \begin{cases} C_{cca} \left(1 - \frac{|\boldsymbol{x}_i - \boldsymbol{x}_j|}{R_{i,A} + R_{j,A}}\right)^2 \frac{\boldsymbol{x}_i - \boldsymbol{x}_j}{|\boldsymbol{x}_i - \boldsymbol{x}_j|} & \text{if } |\boldsymbol{x}_i - \boldsymbol{x}_j| \leq R_{i,A} + R_{j,A}, \\ \mathbf{0} & \text{otherwise,} \end{cases} \tag{6}$$

where the coefficient $C_{cca}$ represents the cell-cell adhesion strength.

On the other hand, the cell-cell repulsion force is activated when two cells start overlapping, so when the distance between the two cell centres $|\boldsymbol{x}_i - \boldsymbol{x}_j|$ is less than the sum of their equivalent radii $R_i + R_j$. This force is used to reproduce the effect of volume exclusion and resistance to cell deformation when a cell is pushed by other cells, and is defined as

$$\boldsymbol{F}_{ccr}^{ij} = \begin{cases} -C_{ccr}\left(1 - \frac{|\boldsymbol{x}_i - \boldsymbol{x}_j|}{R_i + R_j}\right)^2 \frac{\boldsymbol{x}_i - \boldsymbol{x}_j}{|\boldsymbol{x}_i - \boldsymbol{x}_j|} & \text{if } |\boldsymbol{x}_i - \boldsymbol{x}_j| \leq R_i + R_j, \\ \boldsymbol{0} & \text{otherwise,} \end{cases} \tag{7}$$

where $C_{ccr}$ coefficient represent the cell-cell repulsion strength.

##### 1.2.2 Cell motility

The cell motility velocity ($\boldsymbol{v}_{i,\text{mot}}$ in Equation (5)) is a result of cell-ECM adhesion to fibres (contact guidance) combined with random biased migration (random motion combined with chemotaxis). It can be written as:

$$\boldsymbol{v}_{i,\text{mot}} = s_{i,\text{mot}}\, \boldsymbol{d}_{i,\text{mot}}, \tag{8}$$

where $s_{i,\text{mot}}$ is the cell speed and $\boldsymbol{d}_{i,\text{mot}}$ is the motility direction influenced by chemotaxis towards a chemical substrate.

As in Botticelli et al.[2], the cell motility speed $s_{i,\text{mot}}$ represents the magnitude of a cell's velocity due to adhesion to the ECM fibres. This speed is directly proportional to the local ECM density $\rho$, since a denser ECM provides more adhesion sites that the cell can adhere to. Therefore, we define $s_{i,\text{mot}}$ as:

$$s_{i,\text{mot}} = 4\, S_{\max}\, \rho, \tag{9}$$

where $S_{\max}$ is the maximum cell speed. The scaling factor of 4 ensures that the maximum speed that an isolated cell can reach by combining adhesive and repulsive interactions is equal to $S_{\max}$ (see Equation (13)).

##### 1.2.3 Cell-ECM repulsion (or resistance)

As demonstrated in Botticelli et al.[2], a dense 3D matrix, in addition to providing a higher number of adhesion sites, can also impede cell migration by occupying space. Therefore, we introduce a cell-ECM repulsion (or resistance) velocity $\boldsymbol{v}_{i,cmr}$, which is also ECM density-dependent and reduces the cell total speed. The repulsion is maximal when the ECM density $\rho$ is equal to 1 (the space is completely filled by ECM, acting as a wall) and is zero when $\rho$ is equal to zero (no ECM is present). This repulsion force opposes cell-cell forces and motility forces, and is defined as

$$\boldsymbol{v}_{i,cmr} = -\left(\boldsymbol{v}_{i,cca} + \boldsymbol{v}_{i,ccr} + \boldsymbol{v}_{i,\mathrm{mot}}\right)\rho. \tag{10}$$

Combining motility and repulsion, we obtain a non-monotonic total cell speed. Starting from Equation (5) and using Equation (10), assuming no cell-cell interactions ($\boldsymbol{v}_{i,cca} + \boldsymbol{v}_{i,ccr} = \mathbf{0}$), the total cell velocity is the sum of motility and repulsion, which gives us

$$\boldsymbol{v}_i = \boldsymbol{v}_{i,\mathrm{mot}} + \boldsymbol{v}_{i,cmr} \tag{11}$$

$$= \boldsymbol{v}_{i,\mathrm{mot}}(1-\rho). \tag{12}$$

Therefore, substituting in Equation (9), the total cell speed is equal to

$$s_i = 4\,S_{\max}\,\rho(1-\rho). \tag{13}$$

This function reaches its maximum value of $S_{\max}$ when the ECM density $\rho$ is equal to 0.5, making it the optimal density for migration. This behaviour is compatible with experiments that have observed similar non-monotonic migration speed responses to ECM density[8].

#### 1.3 ECM remodelling

In the model, when a cell comes into contact with an ECM voxel, it remodels the matrix substrate by changing the ECM average fibre orientation $\boldsymbol{f}$, average anisotropy $a$, and local density $\rho$ in relation to the cell's movements. Remodelling is applied only to the single voxel at the cell front and is directed along the cell's migration vector.

##### 1.3.1 Orientation

As in Metzcar et al.[11], we assume that the fibre orientation $\boldsymbol{f}$ gets modified by the cell $C_i$ as a result of its active interaction with the fibres, in its direction of movement $\boldsymbol{d}_{i,\mathrm{mot}}$ and proportionally to its speed $s_{i,\mathrm{mot}}$ due to motility. To account for the fact that a fibre is described by a vector but the orientation is physically not relevant, we ensure that the dot product between the fibre orientation $\boldsymbol{d}_{\boldsymbol{f}}$ and the direction of movement $\boldsymbol{d}_{i,\mathrm{mot}}$ is positive. If it is negative, we change the sign of the fibre orientation. This way, we consider the smaller angle between the two vectors and remodel accordingly. The change in orientation is then modelled as follows:

$$\frac{d\boldsymbol{f}}{dt} = -r_f \left( \boldsymbol{f} - \frac{\boldsymbol{d}_{i,\mathrm{mot}}}{||\boldsymbol{d}_{i,\mathrm{mot}}||} \right), \tag{14}$$

where $r_f = r_{f0}\, s_{i,\mathrm{mot}}(1-a)$, with $r_{f0}$ base fibre reorientation rate. The vector $\boldsymbol{f}$ is normalised after each update. Notice that when the anisotropy $a$ is 1, the fibre orientation cannot change. Therefore, we always initialise the simulations with an initial anisotropy $a < 1$.

##### 1.3.2 Anisotropy

The local anisotropy $a$ ranges from 0 to 1, where 0 means no fibre-fibre correlation, and 1 means complete fibre-fibre correlation (the fibres within the voxel are parallel to each other). As in Metzcar et al.[11], for simplicity we assume that the ECM anisotropy always increases and is remodelled proportionally to the motility speed $s_{i,\mathrm{mot}}$ in the following way:

$$\frac{da}{dt} = r_a(1-a), \tag{15}$$

where $r_a = r_{a0}\, s_{i,\mathrm{mot}}$, with $r_{a0}$ base fibre realignment rate.

##### 1.3.3 Density

As in Metzcar et al.[11], the ECM density $\rho$ is degraded according to

$$\frac{d\rho}{dt} = \begin{cases} -r_{d,\mathrm{deg}}\left(\rho - \rho_{\mathrm{target}}\right), & \text{if } \rho > \rho_{\mathrm{target}} \\ 0, & \text{otherwise} \end{cases} \tag{16}$$

with degradation rate $r_{d,\mathrm{deg}}$. In our model, ECM is degraded towards a global target density $\rho_{\mathrm{target}}$. For simplicity, we assume this target density is equal to the intermediate ECM density at which cell migration speed is maximal in our model, corresponding to $\rho = 0.5$ (Equation (13)).

### 2 Parameter values

| Symbol | Parameter | Values | Dimensions |
|---|---|---|---|
| $\Delta t_{mech}$ | Mechanical time step | 0.1 [6] | min |
| $\Delta t_{cell}$ | Phenotype time step | 6 [6] | min |
| $\rho$ | ECM density | [0,1] | Dimensionless |
| $V$ | Maximum cell volume | 2494 [1;6] | $\mu$m$^3$ |
| $R$ | Maximum cell radius | 8.4127 [1;6] | $\mu$m |
| $R_A$ | Maximum interaction radius | $1.25\times R$ [9] | $\mu$m |
| $C_{cca}$ | Cell-cell adhesion strength | 0.4 [9;6] | $\mu$m min$^{-1}$ |
| $C_{ccr}$ | Cell-cell repulsion strength | 10 [9;6] | $\mu$m min$^{-1}$ |
| $S_{max}$ | Maximum cell speed | 0.2 | $\mu$m min$^{-1}$ |
| $D$ | Chemical substrate diffusion coefficient | 100,000 [5] | $\mu$m$^2$ min$^{-1}$ |
| $\lambda$ | Chemical substrate decay rate | 0.1 [12;6;10] | min$^{-1}$ |
| $\bar{S}_i$ | Cell $C_i$'s secretion rate of chemical substrate | 0.0 [11] | min$^{-1}$ |
| $\bar{U}_i$ | Cell $C_i$ uptake rate of chemical substrate | 10 [6] | min$^{-1}$ |
| $\theta$ at $t=0$ | Initial oxygen | 38 [11] | mmHg |
| $\widetilde{\theta}$ | Half-maximum oxygen | 21.5 [13;8] | mmHg |
| - | Hill power | 4 [8] | Dimensionless |

Table 2: List of parameters used in all simulations. All other parameters are set to their default values as used in PhysiCell 1.12.0 [6].

| Symbol | Parameter | Values | Dimensions |
|---|---|---|---|
| $a$ at $t=0$ | Initial anisotropy | 0.0 | Dimensionless |
| $\rho$ at $t=0$ | Initial density | 0.5 | Dimensionless |
| $\alpha$ | ECM sensitivity | [0,1] | Dimensionless |
| $\beta$ | Chemotaxis bias | [0,1] | Dimensionless |
| $r_{f0}$ | Reorientation rate | 0.01 [11] | min$^{-1}$ |
| $r_{a0}$ | Realignment rate | 0.0001 [11] | min$^{-1}$ |
| $r_{d,deg}$ | Degradation rate | 0.0 | min$^{-1}$ |
| $r_{d,disp}$ | Displacement rate | 0.0 | min$^{-1}$ |
| $r_{div}$ | Proliferation rate | 0.0 | min$^{-1}$ |

Table 3: List of parameters used in Section 3.1

| Symbol | Parameter | Values | Dimensions |
|---|---|---|---|
| $a$ at $t=0$ | Initial anisotropy | 0.3, 0.5, 0.7 | Dimensionless |
| $\rho$ at $t=0$ | Initial density | 0.5 | Dimensionless |
| $\alpha$ | ECM sensitivity | [0,1] | Dimensionless |
| $\beta$ | Chemotaxis bias | [0,1] | Dimensionless |
| $r_{f0}$ | Reorientation rate | 0.01 [11] | min$^{-1}$ |
| $r_{a0}$ | Realignment rate | 0.0001 [11] | min$^{-1}$ |
| $r_{d,deg}$ | Degradation rate | 0.0 | min$^{-1}$ |
| $r_{d,disp}$ | Displacement rate | 0.0 | min$^{-1}$ |
| $r_{div}$ | Proliferation rate | 0.0 | min$^{-1}$ |

Table 4: List of parameters used in Section 3.2

| Symbol | Parameter | Values | Dimensions |
|---|---|---|---|
| $a$ at $t = 0$ | Initial anisotropy | 0.0 | Dimensionless |
| $\rho$ at $t = 0$ | Initial density | 0.7 | Dimensionless |
| $\alpha$ | ECM sensitivity | 0.8 | Dimensionless |
| $\beta$ | Chemotaxis bias | 0.2 | Dimensionless |
| $r_{f0}$ | Reorientation rate | 0.01 [11] | $\text{min}^{-1}$ |
| $r_{a0}$ | Realignment rate | 0.0001 [11] | $\text{min}^{-1}$ |
| $r_{d,deg}$ | Degradation rate | [0,1] | $\text{min}^{-1}$ |
| $\rho_{target}$ | Degradation target value | 0.5 | Dimensionless |
| $r_{d,disp}$ | Displacement rate | [0,1] | $\text{min}^{-1}$ |
| $r_{div}$ | Proliferation rate | 0.00072 [6] | $\text{min}^{-1}$ |

Table 5: List of parameters used in Section 3.3

| Symbol | Parameter | Values | Dimensions |
|---|---|---|---|
| $a$ at $t = 0$ | Initial anisotropy | 0.7 | Dimensionless |
| $\rho$ at $t = 0$ | Initial density | 0.7 | Dimensionless |
| $\alpha$ | ECM sensitivity | 0.8 | Dimensionless |
| $\beta$ | Chemotaxis bias | 0.2 | Dimensionless |
| $r_{f0}$ | Reorientation rate | [0,1] | $\text{min}^{-1}$ |
| $r_{a0}$ | Realignment rate | 0.0001 [11] | $\text{min}^{-1}$ |
| $r_{d,deg}$ | Degradation rate | 0.01 | $\text{min}^{-1}$ |
| $\rho_{target}$ | Degradation target value | 0.5 | Dimensionless |
| $r_{d,disp}$ | Displacement rate | 0.1 | $\text{min}^{-1}$ |
| $r_{div}$ | Proliferation rate | 0.00072 [6] | $\text{min}^{-1}$ |

Table 6: List of parameters used in Section 3.4

### References


[1] D. B. Agus, J. F. Alexander, W. Arap, S. Ashili, J. E. Aslan, R. H. Austin, V. Backman, K. J. Bethel, R. Bonneau, W.-C. Chen, C. Chen-Tanyolac, N. C. Choi, S. A. Curley, M. Dallas, D. Damania, P. C. W. Davies, P. Decuzzi, L. Dickinson, L. Estevez-Salmeron, V. Estrella, M. Ferrari, C. Fischbach, J. Foo, S. I. Fraley, C. Frantz, A. Fuhrmann, P. Gascard, R. A. Gatenby, Y. Geng, S. Gerecht, R. J. Gillies, B. Godin, W. M. Grady, A. Greenfield, C. Hemphill, B. L. Hempstead, A. Hielscher, W. D. Hillis, E. C. Holland, A. Ibrahim-Hashim, T. Jacks, R. H. Johnson, A. Joo, J. E. Katz, L. Kelbauskas, C. Kesselman, M. R. King, K. Konstantopoulos, C. M. Kraning-Rush, P. Kuhn, K. Kung, B. Kwee, J. N. Lakins, G. Lambert, D. Liao, J. D. Licht, J. T. Liphardt, L. Liu, M. C. Lloyd, A. Lyubimova, P. Mallick, J. Marko, O. J. T. McCarty, D. R. Meldrum, F. Michor, S. M. Mumenthaler, V. Nandakumar, T. V. O'Halloran, S. Oh, R. Pasqualini, M. J. Paszek, K. G. Philips, C. S. Poultney, K. Rana, C. A. Reinhart-King, R. Ros, G. L. Semenza, P. Senechal, M. L. Shuler, S. Srinivasan, J. R. Staunton, Y. Stypula, H. Subramanian, T. D. Tlsty, G. W. Tormoen, Y. Tseng, A. van Oudenaarden, S. S. Verbridge, J. C. Wan, V. M. Weaver, J. Widom, C. Will, D. Wirtz, J. Wojtkowiak, P.-H. Wu, and The Physical Sciences - Oncology Centers Network. A physical sciences network characterization of non-tumorigenic and metastatic cells. 3(1):1449. ISSN 2045-2322. doi: 10.1038/srep01449. URL https://www.nature.com/articles/srep01449.

[2] M. Botticelli, J. Metzcar, T. Phillips, S. Cox, P. Keshavanarayana, and F. Spill. A hybrid computational model of cancer spheroid growth with ribose-induced collagen stiffening. *Frontiers in Bioengineering and Biotechnology*, 13:1515962, 2025. ISSN 2296-4185. doi: 10.3389/fbioe.2025.1515962.

[3] L. Capuana, A. Boström, and S. Etienne-Manneville. Multicellular scale front-to-rear polarity in collective migration. *Current Opinion in Cell Biology*, 62:114–122, 2020. ISSN 09550674. doi: 10.1016/j.ceb.2019.10.001.

[4] D. Drasdo, R. Kree, and J. S. McCaskill. Monte carlo approach to tissue-cell populations. *Physical Review E*, 52(6):6635–6657, 1995. ISSN 1063-651X, 1095-3787. doi: 10.1103/PhysRevE.52.6635.

[5] A. Ghaffarizadeh, S. H. Friedman, and P. Macklin. BioFVM: an efficient, parallelized

diffusive transport solver for 3-D biological simulations. *Bioinformatics*, 32(8):1256–1258, 2016. ISSN 1367-4803, 1460-2059. doi: 10.1093/bioinformatics/btv730.

[12] M. R. Owen, H. M. Byrne, and C. E. Lewis. Mathematical modelling of the use of macrophages as vehicles for drug delivery to hypoxic tumour sites. *Journal of Theoretical Bi-

*ology*, 226(4):377–391, 2004. ISSN 0022-5193. doi: https://doi.org/10.1016/j.jtbi.2003.09.004. URL https://www.sciencedirect.com/science/article/pii/S0022519303003412.

[13] H. L. Rocha, I. Godet, F. Kurtoglu, J. Metzcar, K. Konstantinopoulos, S. Bhoyar, D. M. Gilkes, and P. Macklin. A persistent invasive phenotype in post-hypoxic tumor cells is revealed by fate mapping and computational modeling. *iScience*, 24(9):102935, 2021. ISSN 25890042. doi: 10.1016/j.isci.2021.102935.
