## Supplementary figures and images for "The role of contact guidance and ECM remodelling in cancer invasion: a computational study"

### figureS1

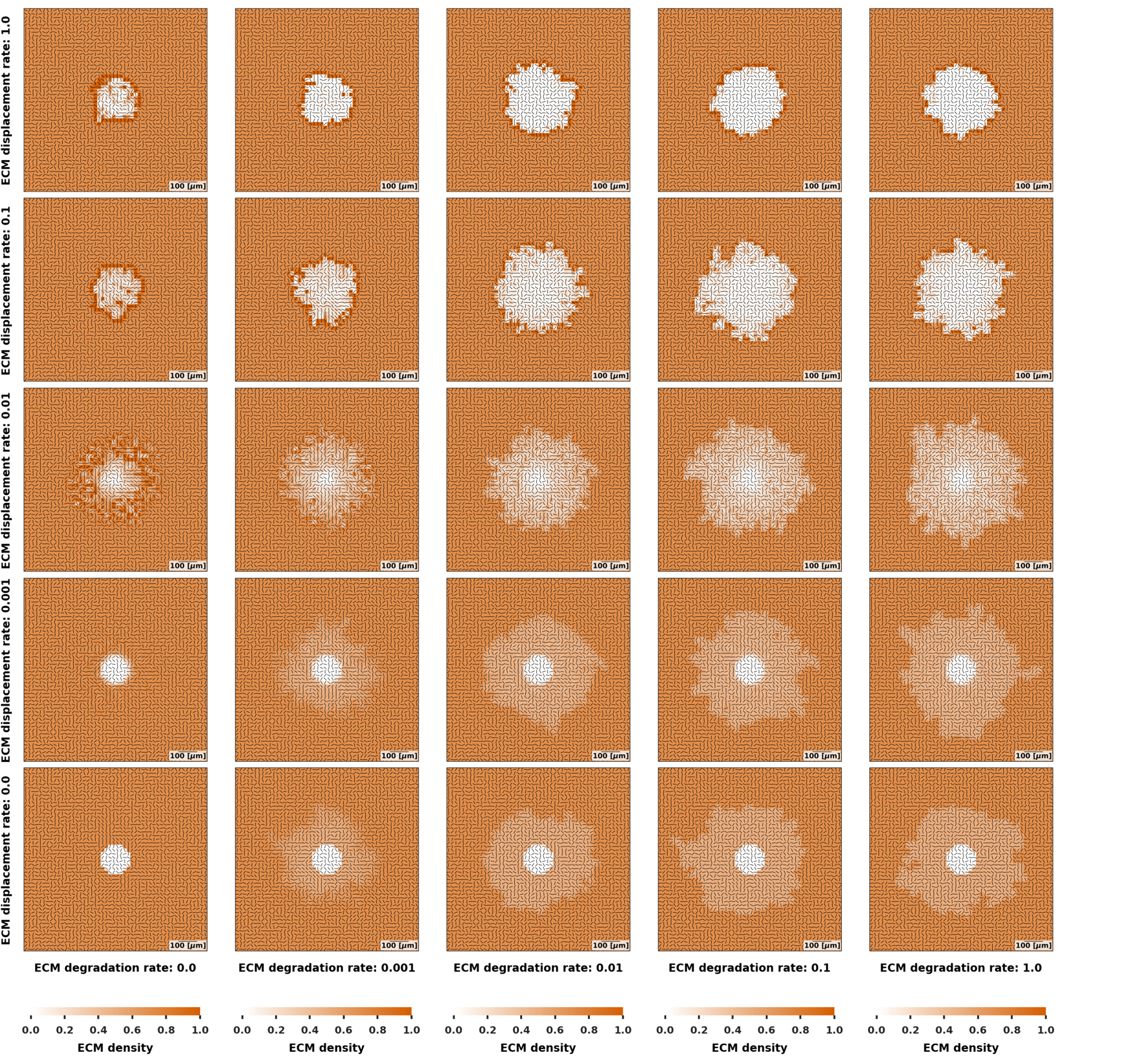

### figureS2

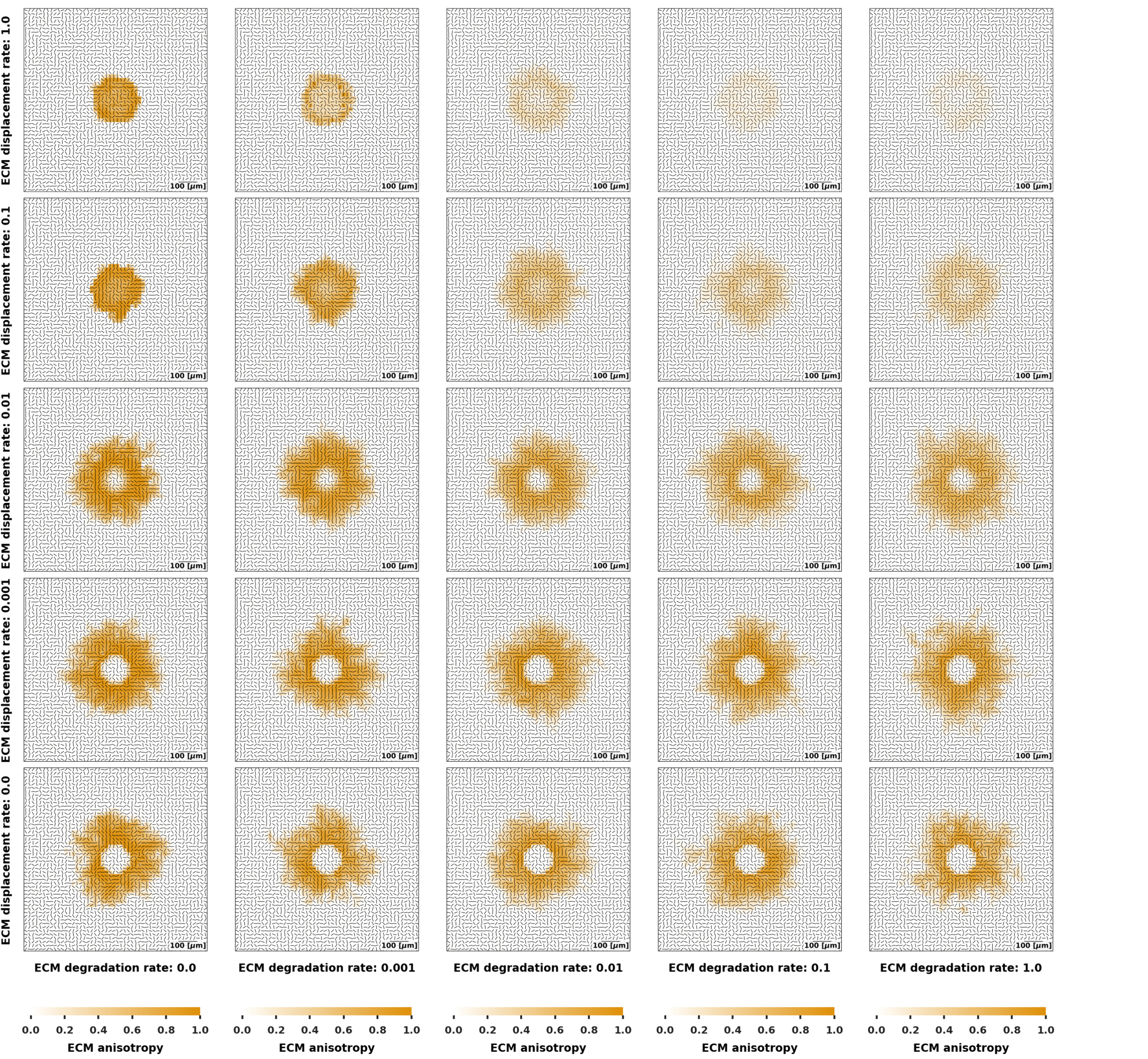

### figureS3

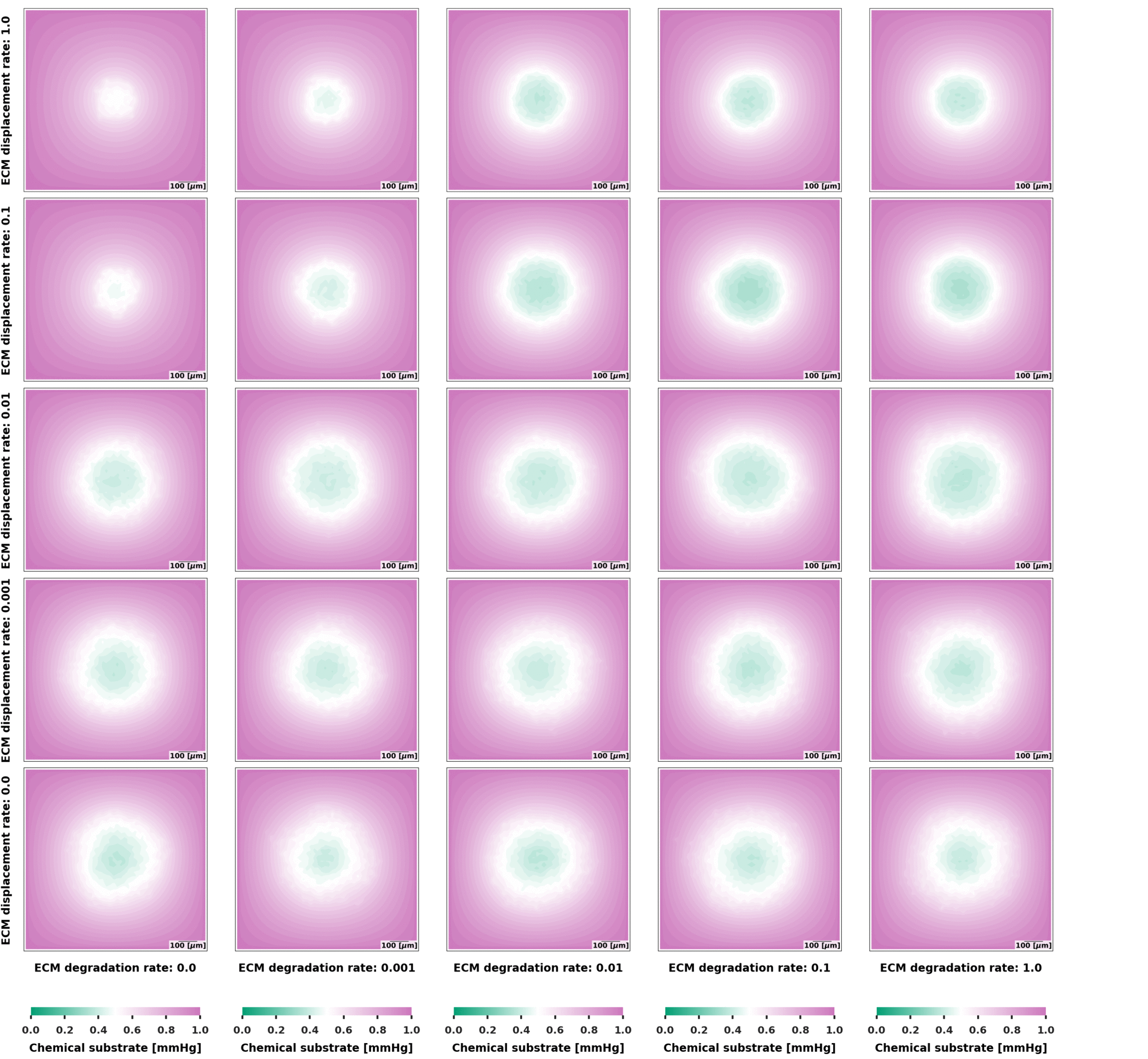

### figureS4

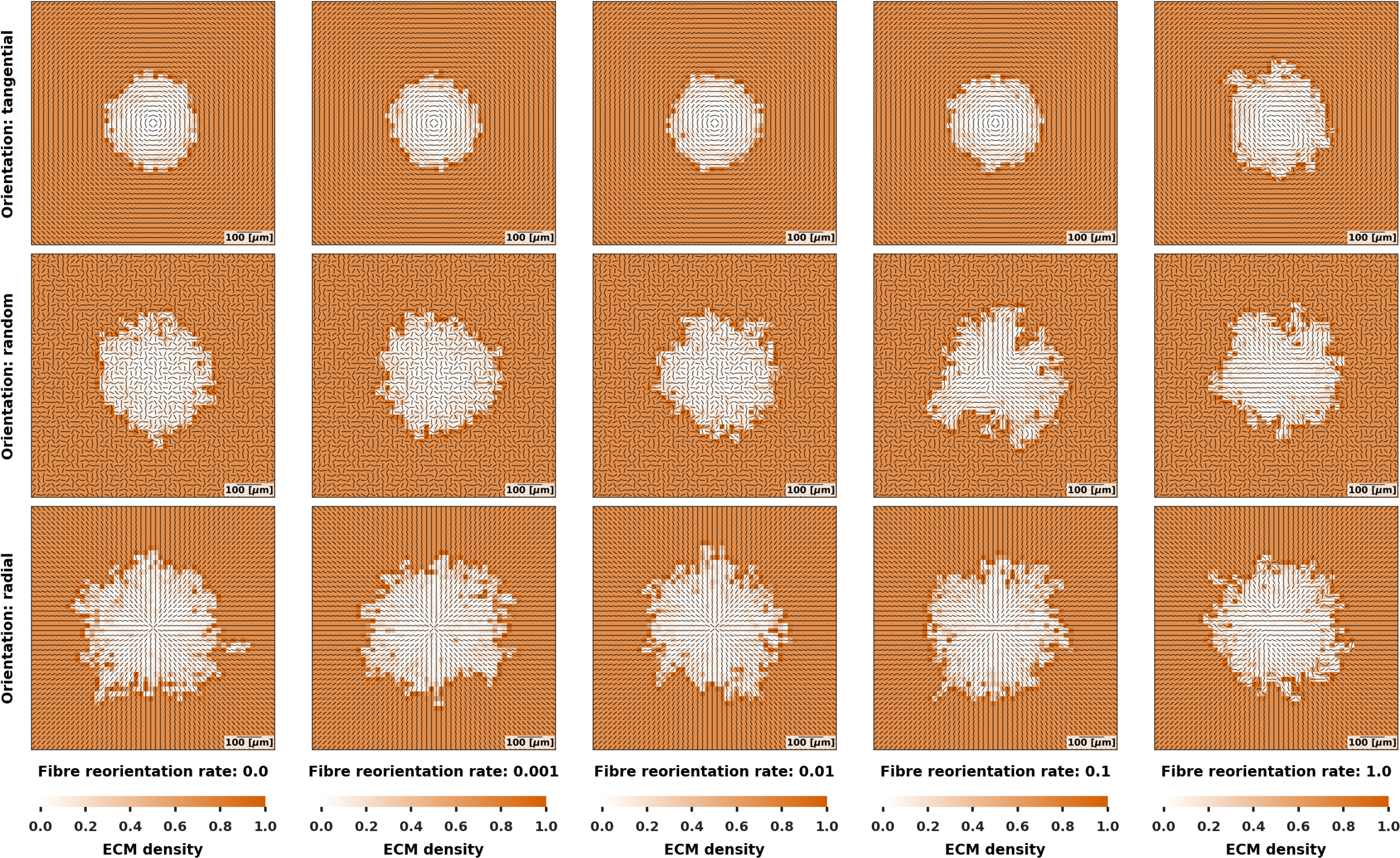

### figureS5

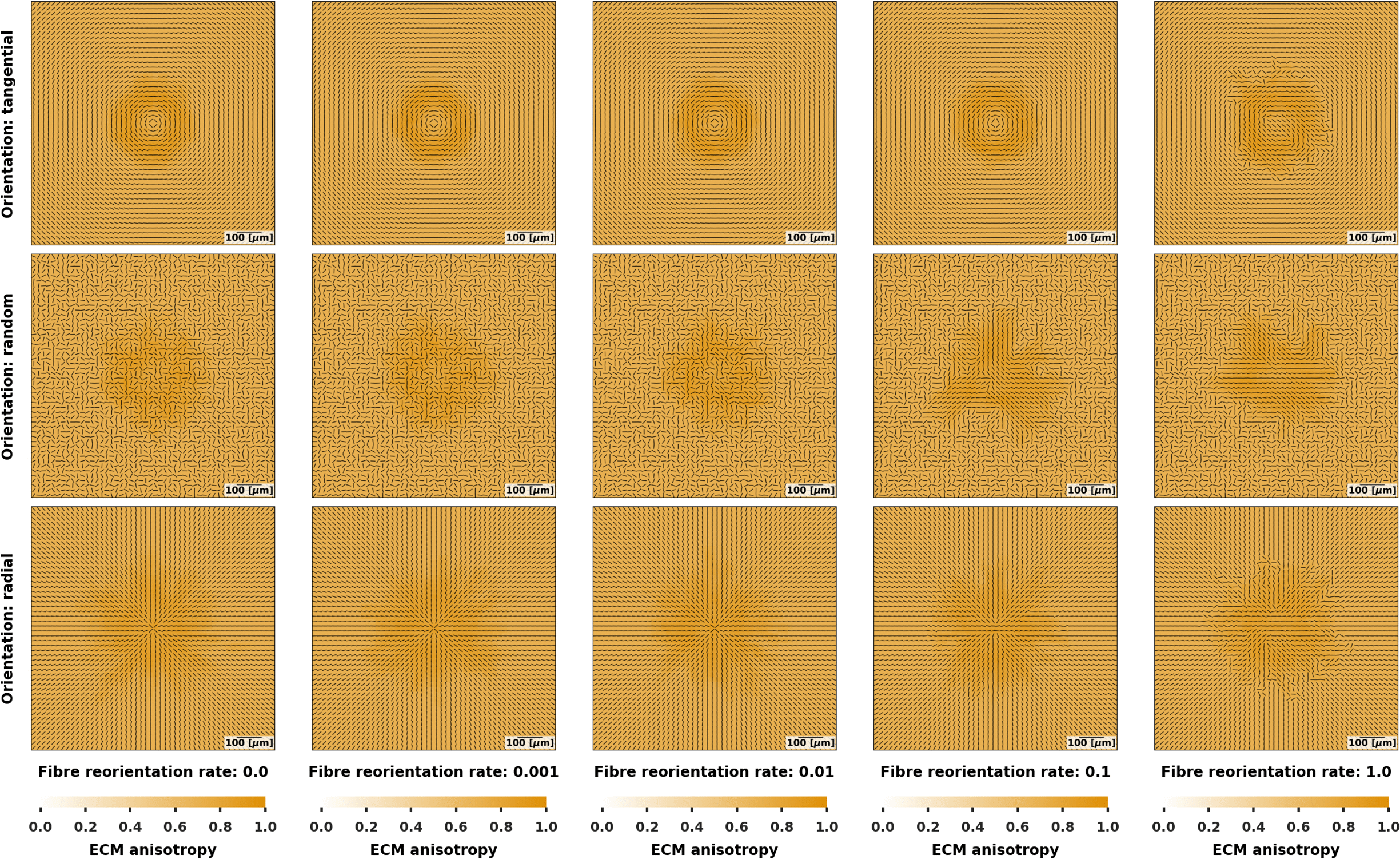

### figureS6

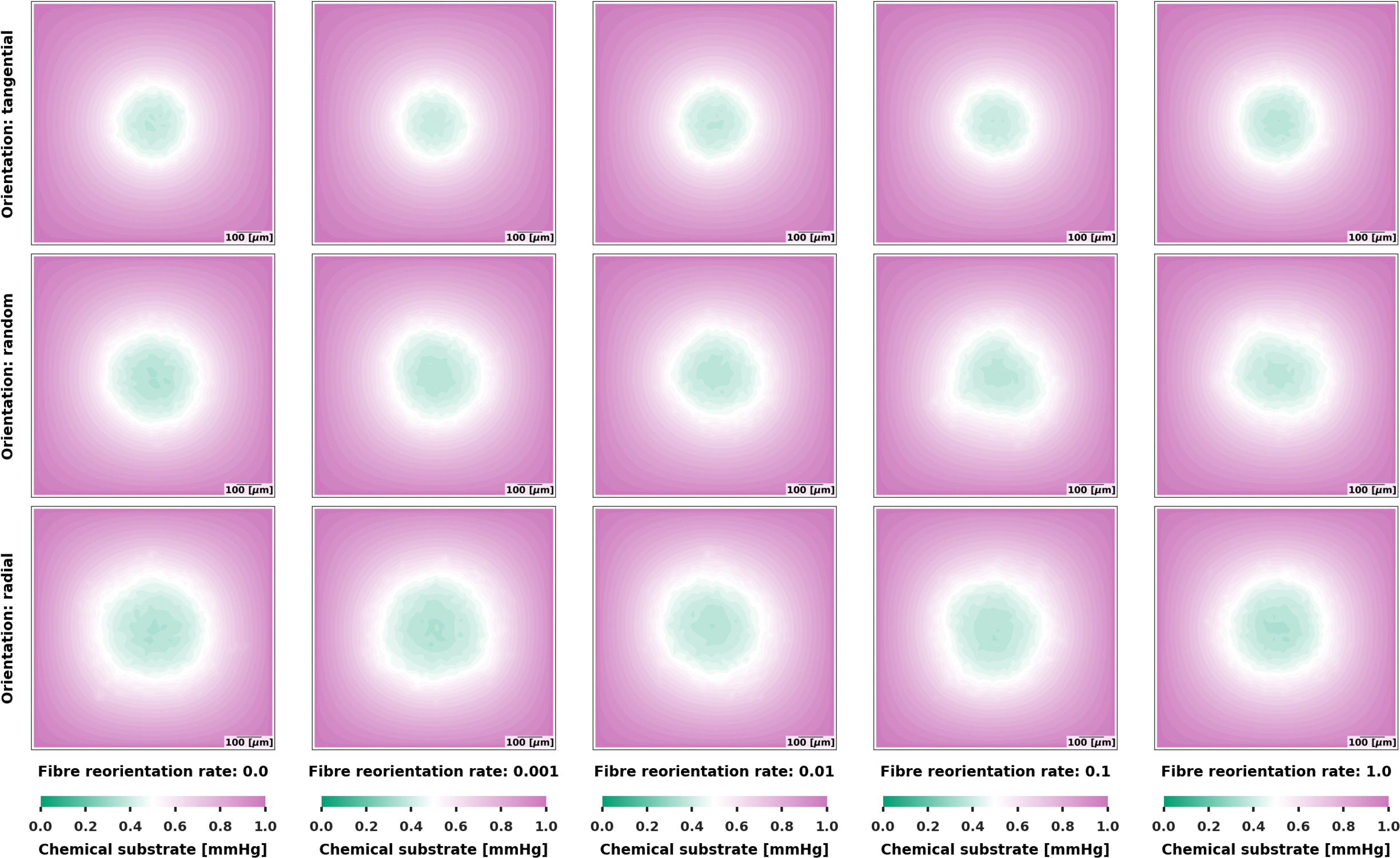
